# Proton-transporting heliorhodopsins from marine giant viruses

**DOI:** 10.1101/2022.03.24.485645

**Authors:** Shoko Hososhima, Ritsu Mizutori, Rei Abe-Yoshizumi, Andrey Rozenberg, Shunta Shigemura, Alina Pushkarev, Masae Konno, Kota Katayama, Keiichi Inoue, Satoshi P. Tsunoda, Oded Béjà, Hideki Kandori

## Abstract

Rhodopsins convert light into signals and energy in animals and microbes. Heliorhodopsins (HeRs), a recently discovered new rhodopsin family, are widely present in archaea, bacteria, unicellular eukaryotes, and giant viruses, but their function remains unknown. Here we report that a viral HeR from *Emiliania huxleyi* virus 202 (V2HeR3) is a light-gated proton channel. V2HeR3 absorbs blue-green lights, and the active intermediate contains the deprotonated retinal Schiff base. Site-directed mutagenesis study revealed that E191 in TM6 constitutes the gate together with the retinal Schiff base. E205 and E215 form a proton accepting group of the Schiff base, whose mutations converted the protein into an outward proton pump. Three environmental viral HeRs from the same group, as well as a more distantly related HeR exhibited similar proton-transport activity, indicating that HeR functions might be diverse similarly to type-1 microbial rhodopsins. Some strains of *E. huxleyi* contain one HeR that is related to the viral HeRs, while its viruses *Eh*V-201 and *Eh*V-202 contain two and three HeRs, respectively. Except for V2HeR3 from *Eh*V-202, none of these proteins exhibit ion-transport activity. Thus, when expressed in the *E. huxleyi* cell membranes, only V2HeR3 has the potential to depolarize the host cells by light, possibly to overcome the host defense mechanisms or to prevent superinfection. The neuronal activity generated by V2HeR3 suggests that it can potentially be used as an optogenetic tools, like type-1 microbial rhodopsins.

## Introduction

Many organisms sense sunlight using rhodopsins (Ernst et al., 2014; Govorunova et al., 2017; Grote et al., 2014; Rozenberg et al., 2021), which are integral membrane proteins that bind retinal chromophores. Rhodopsins are classified into type-1 microbial or type-2 animal rhodopsins, which contain all-*trans* or 11-*cis* retinal, respectively, as the chromophore. Type-2 animal rhodopsins function as G-protein coupled receptors, while the functions of type-1 microbial rhodopsins are highly diverse and include light-driven ion pumps, light-gated ion channels, light sensors, and light-activated enzymes. Ion-transporting rhodopsins are used as the main tools in optogenetics (Deisseroth and Hegemann, 2017). In addition to type-1 and type-2 rhodopsins, a previously unrecognised diverse family, heliorhodopsins (HeRs), was recently discovered using functional metagenomics (Pushkarev et al., 2018). Although HeRs are phylogenetically distant from the type-1 rhodopsins, structures and photocycles of HeRs resemble those of type-1 rhodopsins (Kovalev et al., 2020; Lu et al., 2020; Pushkarev et al., 2018; Shihoya et al., 2019). The most noticeable difference is that HeRs have inverted membrane topology compared to type-1 and -2 rhodopsins (Pushkarev et al., 2018; Shihoya et al., 2019).

Similarly to type-1 rhodopsins, HeRs are encoded in genomes of archaea, bacteria, unicellular eukaryotes, and giant viruses. Physiological functions of HeRs, however, remain unknown. Previous studies did not detect any ion-transport activity in HeRs, and from their long-photocycle, a sensory function was suggested (Pushkarev et al., 2018; Shihoya et al., 2019). Analogously, no ion-transport function is consistent with the crystal structures of HeRs from Thermoplasmatale1s archaeon (*Ta*HeR) (Shihoya et al., 2019) and actinobacterial 48C12 (Kovalev et al., 2020; Lu et al., 2020), where the interior of the extracellular half is highly hydrophobic. It should be noted however that structural information is limited to only a few HeRs from archaea and bacteria. HeRs are a highly diverse group (Kovalev et al., 2020) for which a variety of functions might be expected. In the present study, we report ion-transport activity for a viral HeR, hinting at an unanticipated diversity of HeRs. *Emiliania huxleyi* (= *Gephyrocapsa huxleyi*) is a globally important marine microalga. Massive blooms of *E. huxleyi* are observable from satellites and have an impact on Earth’s climate (Tyrrell and Merico, 2004). Giant double-stranded DNA viruses from the genus *Coccolithovirus* (Phycodnaviridae) are known to infect *E. huxleyi* and collapse its blooms (Bidle et al., 2007; Vardi et al., 2009). Curiously, *coccolithoviruses* encode HeRs in their genomes, with some isolates, such as *Eh*V-202, having up to three HeR genes (Fig. 1A). One of the HeR genes from *Eh*V-202 (AET42570.1; V2HeR2) has been expressed before and failed to demonstrate any light-dependent ion transporting activity (Shihoya, 2019). In contrast, here we detected photocurrents for another HeR from *Eh*V-202. Molecular mechanism and functional role will be discussed for ion-transporting HeRs from marine giant viruses.

**Figure 1.**
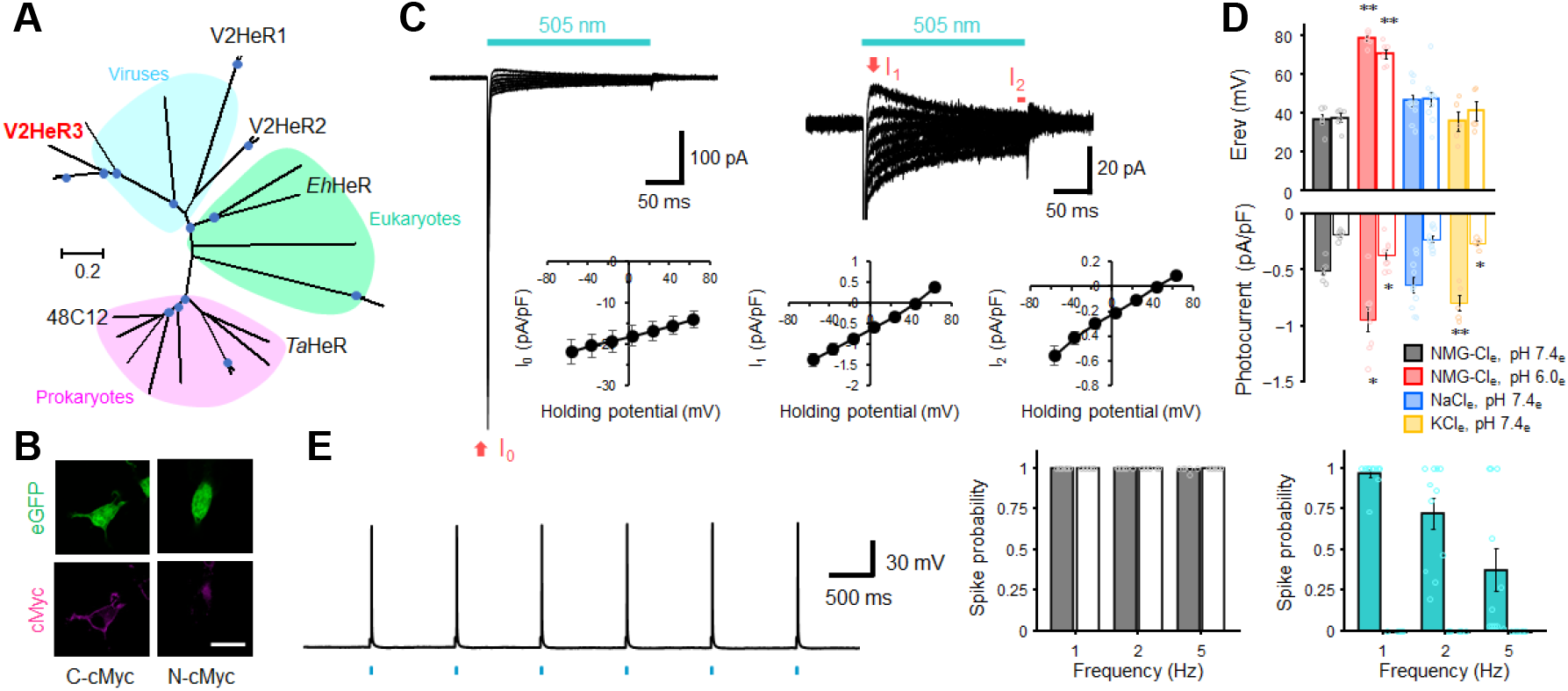
Light-gated inward proton transport of a viral heliorhodopsin from *Emiliania huxleyi* virus 202 (V2HeR3). (A) Phylogenetic tree of heliorhodopsins (HeRs), which includes three viral HeRs from *E. huxleyi* 202 (V2HeR1-3), a eukaryotic HeR from *E. huxleyi* (Ehux-HeR), an archaeal HeR (*Ta*HeR) and a bacterial HeR (48C12). (B) eGFP fluorescence (top, green) and immunofluorescence staining (bottom, magenta) observation of V2HeR3 with a cMyc epitope tag at the C terminus (left) and the N terminus (right) in cultured ND7/23 cells. Scale bar, 20 μm. (C) Electrophysiological measurements of V2HeR3-driven photocurrent in ND7/23 cells. The cells were illuminated with light (λ = 505 nm, 24.5 mW/mm^2^) during the time region shown by the blue bars. The membrane voltage was clamped from −60 to +60 mV for every 20-mV step. The pipette solution was 110 mM NMG-Cl, pHi 7.4, the bath solution was 140 mM NaCl, pHe 7.4. n = 10 cells. (D) Corresponding reversal voltage (E_rev_) for each internal condition (upper), and comparison of photocurrent amplitudes at 0 mV for different internal cations (bottom). Square-block bar graph indicates E_rev_ or amplitude from peak photocurrent, open bar graphs indicate E_rev_ or amplitude from steady-state photocurrent. The pipette solution was 110 mM NMG-Cl, pHi 7.4, the bath solution was 140 mM NMG-Cl, pHe 7.4 (black), 140 mM NMG-Cl, pHe 6.0 (red), 140 mM NaCl, pHe 7.4 (blue) or 140 mM KCl, pHe 7.4 (green). n = 5 to 10 cells. (*p<0.05, ** p<0.01). (E) Sample responses of a V2HeR3-expressing neuron to 10 ms light pulses (left, λ = 505 nm, 24.5 mW/mm^2^) at 10 Hz. Comparison of spike probability by electrical stimulation (middle, 300 pA currents injections) or light stimulation (light, λ = 505 nm, 24.5 mW/mm^2^). The Square-block bar indicates spike probability from V2HeR3-expressing neurons, the open bar indicates spike probability from the neurons without V2HeR3. n = 6 to 11 cells.

## Results

We first targeted one HeR from *Eh*V-202 (AET42421.1; V2HeR3), and applied patch-clamp recording after expressing it in mammalian cells (ND7/23). We visualised protein expression in ND7/23 cells by a tagged-eGFP and a cMyc-epitope tag. Although membrane localization of the expressed proteins was weak (eGFP signals in Fig. 1B), clear fluorescence from anti-cMyc tag was observed only from the C-terminal cMyc, but not from the N-terminal cMyc (magenta in Fig. 1B), indicating that the C-terminus faces the extracellular side, similarly to other HeRs (Pushkarev et al., 2018; Shihoya et al., 2019). Photocurrents in Fig. 1C exhibit a sharp negative transient peak (I_0_), which rapidly drops into a relatively broad peak component (I_1_), followed by a steady-state current component (I_2_) during the illumination. Although I_0_ does not necessarily indicate ion transport, the presence of steady-state currents is an unequivocal indication of ion transport which is maintained for a long time span (Fig. S1). Photocurrent amplitude increases linearly with light intensity (Fig. S2), suggesting photocurrents owing to single-photon events. From the current-voltage (I-V) plot in Fig. 1c, it can be seen that I_1_ and I_2_ exhibit a steep and linear voltage dependency with a reversal potential (E_rev_) at +30 mV and +40 mV respectively, while I_0_ is always negatively directed and exhibits a weak voltage dependency.

Voltage-dependent positive and negative currents of I_1_ and I_2_ (Fig. 1C) imply a light-gated channel function. The I-V plot under the symmetric ionic conditions on both sides of the cells without metal cation (pH 7.4) was almost linear with E_rev_ of about +40 mV (Fig. S3). Lowering the extracellular pH from 7.4 to 6.0 resulted in an E_rev_ shift from +40 mV to +70 mV (Fig. 1D), which was also observed when pH_i_ was lowered (Fig. S3). In contrast, replacing the solutions with Na^+^ or K^+^ did not show any significant E_rev_ shift, suggesting that V2HeR3 is a light-gated proton channel. It should be noted that channelrhodopsin 2 from *Chlamydomonas reinhardtii* (ChR2) (Nagel et al., 2003), a standard depolarization tool in optogenetics (Boyden et al., 2005; Deisseroth and Hegemann, 2017; Ishizuka et al., 2006), possesses an outward proton pump activity (Feldbauer et al., 2009). Significant positive E_rev_ suggests that V2HeR3 also possesses the activity of a proton pump, though the direction is inward. E_rev_ was also altered dependent on the extracellular monovalent anions, but independent on the intracellular side (Fig. S4). This suggests that V2HeR3 contains a binding site for a monovalent anion at the extracellular side affecting proton transport. Illumination of yeast cells (*P. pastoris*) expressing V2HeR3 increased solution pH, which was abolished by the addition of the proton uncoupler CCCP (Fig. S5). This result supports V2HeR3 transporting solely protons.

To test the applicability of V2HeR3 for optogenetic manipulation of neuronal activity, we next expressed V2HeR3 in cultured cortical neurons. Short light pulses successfully triggered action potentials in the transfected cells at 1 Hz (Fig. 1E). Spike probability reached close to 1.0 at 1 Hz illumination whereas it decreased when illuminated at 2 and 5 Hz. The light power required for inducing the action potential was between 2-16 mW/mm_2_, in the same range as required for ChR2 (Fig. S6) (Boyden et al., 2005; Ishizuka et al., 2006). The neurons are excitable by current injections at each frequency even after the V2HeR3 expression, confirming that V2HeR3 itself does not harm the neurons (Fig. 1E). These results indicate that V2HeR3 can be used for optical manipulation of neuronal excitability, although the frequency is limited to up to ca. 1 Hz.

Molecular properties of V2HeR3 were studied for the purified protein heterologously expressed in *P. pastoris* cells. The purified sample demonstrated an absorption maximum at 500 nm (Fig. 2A). HPLC analysis revealed that V2HeR contains 64% all-*trans* retinal in the dark, which was converted to the 13-*cis* form by light (Fig. S7). The pKa of the Schiff base and its counterion were 14.9 and 4.3, respectively (Fig. S8), being close to those of *Ta*HeR and 48C12 (Pushkarev et al., 2018; Shihoya et al., 2019). Although the photocycle upon illumination was slow as for other HeRs, the M intermediate was long-lived and directly returned to the original state (Fig. 2B). Formation of the M intermediate possessed μs and ms components, suggesting that the M intermediate is the conducting state of the light-gated proton channel (Fig. 2C). We then studied photointermediate states at low temperatures. Fig. 2D shows formations of the K and M intermediate at 100 and 250 K, respectively, and their photoequilibria with the unphotolyzed state. FTIR analysis at 77 K (Fig. S9) revealed a peak pair at 1200 (−)/1193 (+) cm^-1^, characteristic of the all-*trans* to 13-*cis* photoisomerization (Fig. 2E). Similar bands were observed for the M intermediate (Fig. 2E and Fig.S10), although the M intermediate generally lacked positive signals. This may originate from (i) vibrations other than retinal, (ii) protonated photointermediates, or (iii) photointermediates of the 13-*cis* photocycle. The negative peak at 1657 cm^-1^ in Fig. 2F is the characteristic vibration of helical amide-I, indicating that structural perturbation α-helix is observed upon retinal isomerization (K intermediate), which is maintained in the M intermediate. Nevertheless, different structural changes in the protein are suggested by the stronger positive peaks at 1628 and 1646 cm^-1^ for K and M intermediates, respectively. A peak pair at 1720 (−)/1712 (+) cm^-1^ is indicative of a hydrogen-bonding change in a protonated carboxylic acid upon retinal isomerization, while the M intermediate shows peaks at 1734 (+)/1720 (−)/1695 (+) cm^-1^ (Fig. 2F). *Ta*HeR and 48C12 showed no spectral changes in this frequency region (Pushkarev et al., 2018; Shihoya et al., 2019), and thus a protonated carboxylic acid of the C=O stretch at 1720 cm^-1^ is unique for V2HeR3.

**Figure 2.**
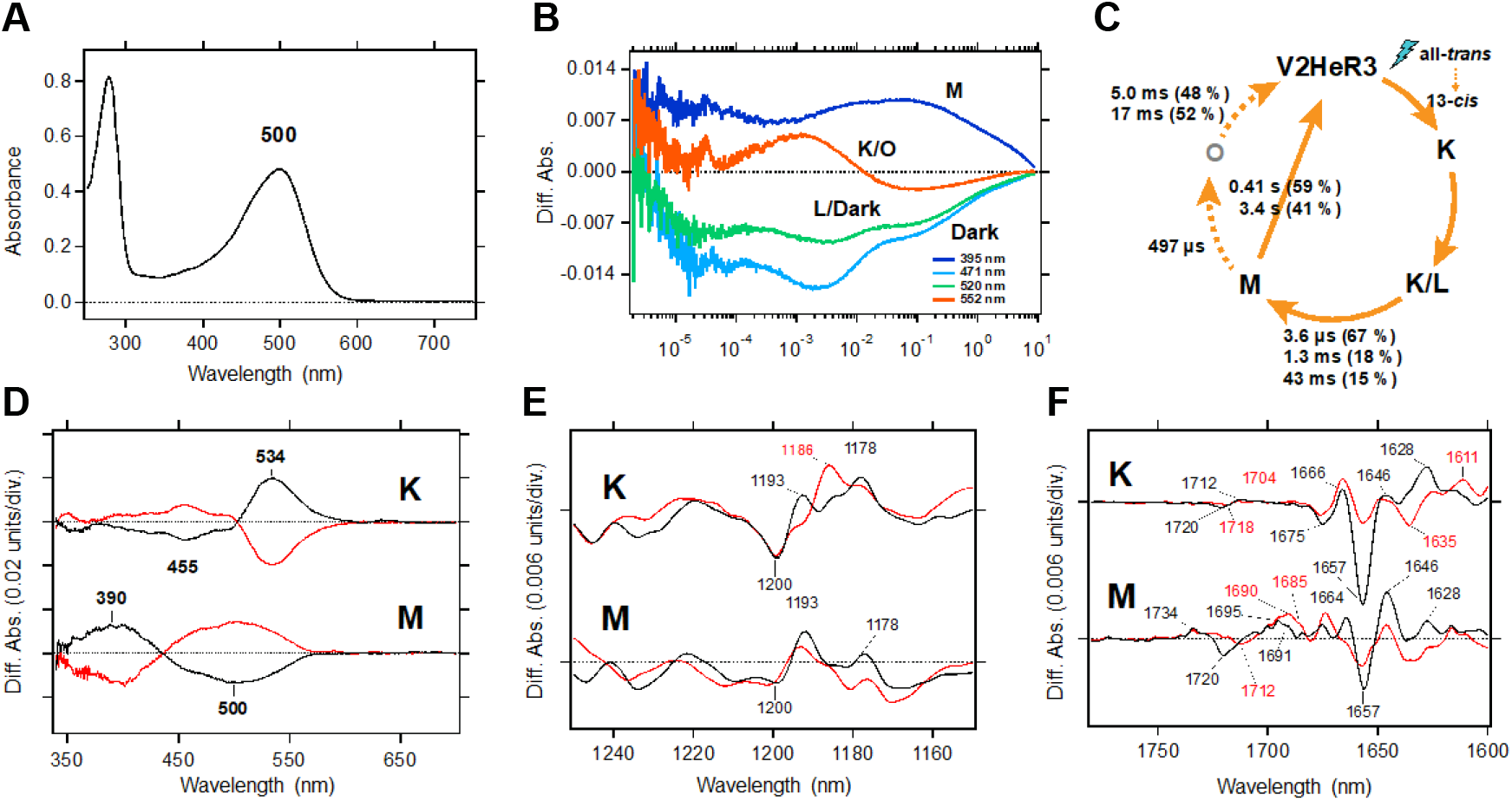
Molecular properties of the purified V2HeR3 proteins expressed in *Pichia pastoris* cells. (A) UV-visible absorption spectrum of V2HeR3 in detergent (0.1 % DDM). (B) Time evolutions of transient absorption changes at characteristic wavelengths of specific photointermediates of V2HeR3. (C) Photocycle of V2HeR3 determined by analyzing the time evolution with multiexponential functions. (D) Light-induced low-temperature K-minus-dark (top) and M-minus-dark (bottom) difference UV-visible spectra of V2HeR3 obtained at 100 and 230 K, respectively. Black curves represent the formation of the K and M intermediates by illuminating at 500 nm and >490 nm, respectively, while red curves represent the reversion from the intermediates by illuminating at >530 nm and 400 nm, respectively. (E) Light-induced low-temperature K-minus-dark (top) and M-minus-dark (bottom) difference FTIR spectra of V2HeR3 obtained at 100 and 230 K, respectively, in the 1250-1150 cm^-1^ region. (F) Light-induced low-temperature K-minus-dark (top) and M-minus-dark (bottom) difference FTIR spectra of V2HeR3 in H_2_O (black) and D_2_O (red) obtained at 100 and 230 K, respectively, in the 1780-1600 cm^-1^ region.

Mechanism of the light-gated proton-transport of V2HeR3 was studied by the use of site-directed mutagenesis. It is well known that internal carboxylates play an important role in ion-transport rhodopsins such as ChR2 and a light-driven proton pump bacteriorhodopsin (BR) (Gerwert et al., 2014; Kandori, 2020; Lórenz-Fonfría and Heberle, 2014). V2HeR3 contains 13 carboxylates, among which D2 and E51/E53 are located at the N-terminus and extracellular loop, respectively, based on the topology prediction, excluding the possibility that these residues are involved in the proton transport (Fig. 3A). We thus prepared 10 mutants affecting the remaining carboxylate positions in all seven helices, and the photocurrents of the D-to-N or E-to-Q mutants were measured. Fig. S11 and S12 show the photocurrents and their I-V plots, respectively, measured by expressing each mutant in ND7/23 cells. Absorption spectra of each mutant in ND7/23 and *P. pastoris* cells were obtained without purification using the hydroxylamine bleaching method (Fig. S13, S14 and S15). Among the 10 mutants, those that affected carboxylates on TM1, 2, 4, and 5 exhibited similar reversal potentials to that of the wild-type (Fig. S11), whereas mutations of E105, E191, E205, E215, and E236 on TM3, TM6 and TM7 were unique (Fig. 3B). E105Q and E191Q in particular diminished photocurrent entirely. E105 is the Schiff base counterion, and its neutralization led to loss of visible absorption (Fig. 3B). This was not the case for 48C12, as the same mutation exhibited visible absorption upon binding chloride (Singh et al., 2019). In the case of E191Q, visible absorption did appear, but without photocurrent. The importance of carboxylate at position 191 is demonstrated further by the fact that the E191D mutant retained ion-transporting activity. Nevertheless, the I-V plots suggest that the E191D mutation diminished the proton channel mode converting the protein entirely to an inward proton pump, as indicated by the negative photocurrents. Positions homologous to E191 are I199 and T178 in *Ta*HeR and BR, respectively, which sandwich F203 and W182, respectively, with retinal (Fig. 3C). It should be noted that W182 exhibits unique conformational changes important for the proton pumping activity in BR (Nango et al., 2016; Subramaniam and Henderson, 2000; Weinert et al., 2019), and that the highly conserved tryptophan in type-1 microbial rhodopsins is replaced with phenylalanine in most HeRs. Interestingly, the proton-transporting V2HeR3 contains tryptophan at this position, which possibly constitutes the gate for ion transport together with E191. In Fig. 3B, E205Q and E215Q show positive photocurrent signals at all the membrane voltages tested (−60 - +60 mV), suggesting proton conduction in the opposite direction (outward proton pump). It is likely that E205 and E215 are the key residues in defining the direction of proton transport. The current shape and the I-V plot of E236Q are similar to those of the wild type, although a shift of E_rev_ to the more positive voltage from +30 mV to +60 mV is observed.

**Figure 3.**
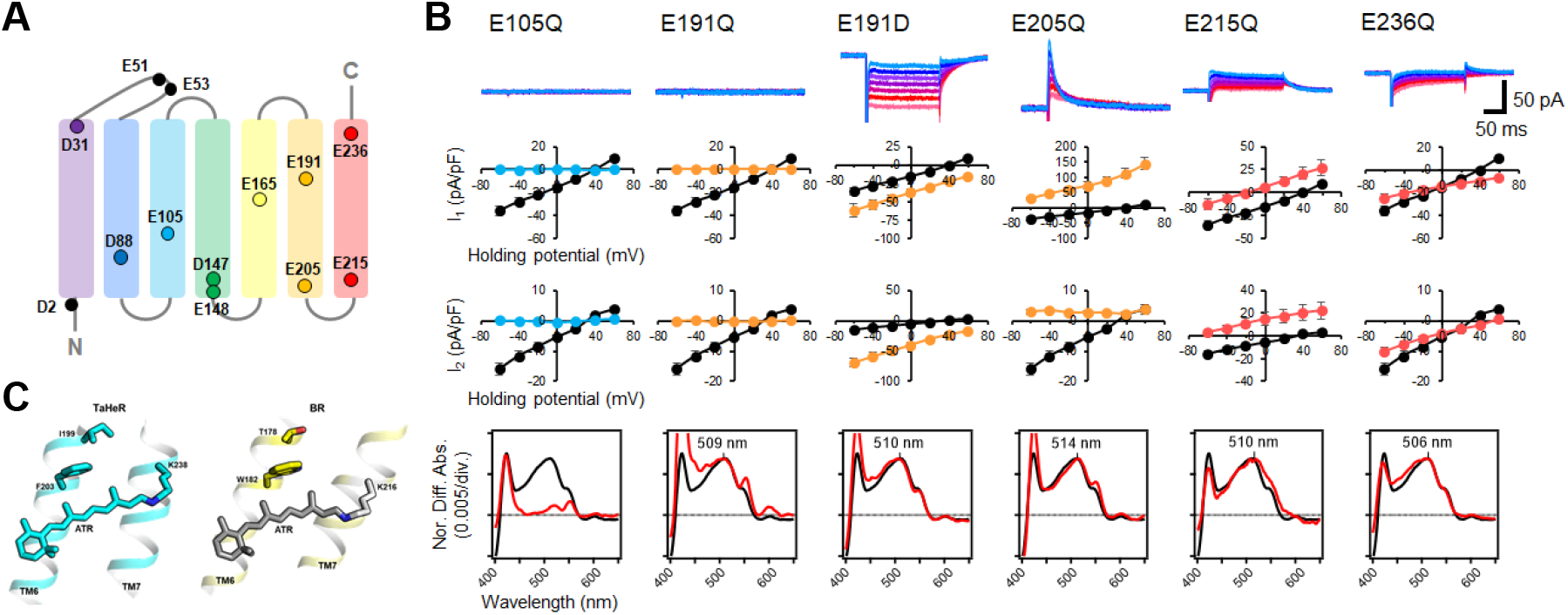
Electrophysiological analysis of the carboxylate mutants of V2HeR3. (A) Aspartate (D) or glutamate (E) in V2HeR3. Among 13 residues, 10 transmembrane aspartate and glutamate were replaced with asparagine (D-to-N) and glutamine (E-to-Q), respectively. (B) Photocurrents (top), I-V plots (middle) and absorption spectra (bottom) of mutants which differ from those of the wild type. Black curves in I-V plots and absorption spectra are the results of the wild type. The pipette solution was 110 mM NaCl, pHi 7.4, the bath solution was 140 mM NaCl, pHe 7.4. n = 6 to 7 cells. (C) Crystal structures of *Ta*HeR (left) and a light-driven proton pump bacteriorhodopsin (BR) (right). Corresponding residues of E191 in V2HeR3 are I199 in *Ta*HeR and T178 in BR. Highly conserved phenylalanine and tryptophan exist in HeRs and type-1 rhodopsins, respectively, while V2HeR3 contains tryptophan at this position.

Despite significant sequence divergence (see Table S1), HeRs from *coccolithoviruses* form a monophylum close to HeRs from haptophytes and other algae, including a HeR gene found in *E. huxleyi* (Fig. S16). Thus, in an attempt to trace the origins of the ion-transporting activity of V2HeR3, we investigated other viral HeRs, as well as the HeR gene from the host species. In addition to V2HeR3, *Eh*V-202 possesses HeRs from two more distantly related clades, V2HeR1 and V2HeR2, that alone are more widespread among *Eh*V isolates (Fig. S17). Nevertheless, none of the four tested HeRs from clades V2HeR1 and V2HeR2 exhibited steady-state photocurrents as demonstrated in Fig. S18 and Fig. 4A. A similarly negative result was obtained for *Eh*HeR, the HeR from the host alga. We then studied a collection of metagenomic viral HeR genes that were closely related to V2HeR3, as well as the more distantly related HeR from *Eh*V-PS401. Fig. S19 and Fig. 4B clearly show that all of them exhibit ion transport activity similar to that of V2HeR3. The I-V plots of VPS401HeR and V*Tara*8957HeR show that the E_rev_ is close to 0 mV, indicating that the proton channel mode is dominant (Fig. 4B). In contrast, ion-transport properties of V*Tara*5482HeR and V*Tara*4616HeR are very similar to that of V2HeR3 judging from their I-V plots, indicating that the proton pump mode is prominent. The absorption spectra of these proteins obtained by hydroxylamine bleaching are shown in Fig. S20. A sequence comparison of the HeRs is shown in Fig. S21, with the key amino acids shown in Fig. 4c. E105 of V2HeR3 is the Schiff base counterion, and E215 is conserved among HeRs. Among the other carboxylate residues, E205 and E236 are not fully conserved among ion-transporting HeRs, while E191 appears to be their hallmark. Interestingly, although relatively uncommon among eukaryotic HeRs, *Eh*HeR that did not demonstrate ion transport, as well as the more distantly-related *Mc*HeR that was tested before (Shihoya et al., 2019), also contain the conserved glutamate E191 (see Fig. 4C and Fig. S16). This indicates that E191 might be essential but not sufficient for ion transport, and that other residues, such as W195 and E/Q205 are required as well.

**Figure 4.**
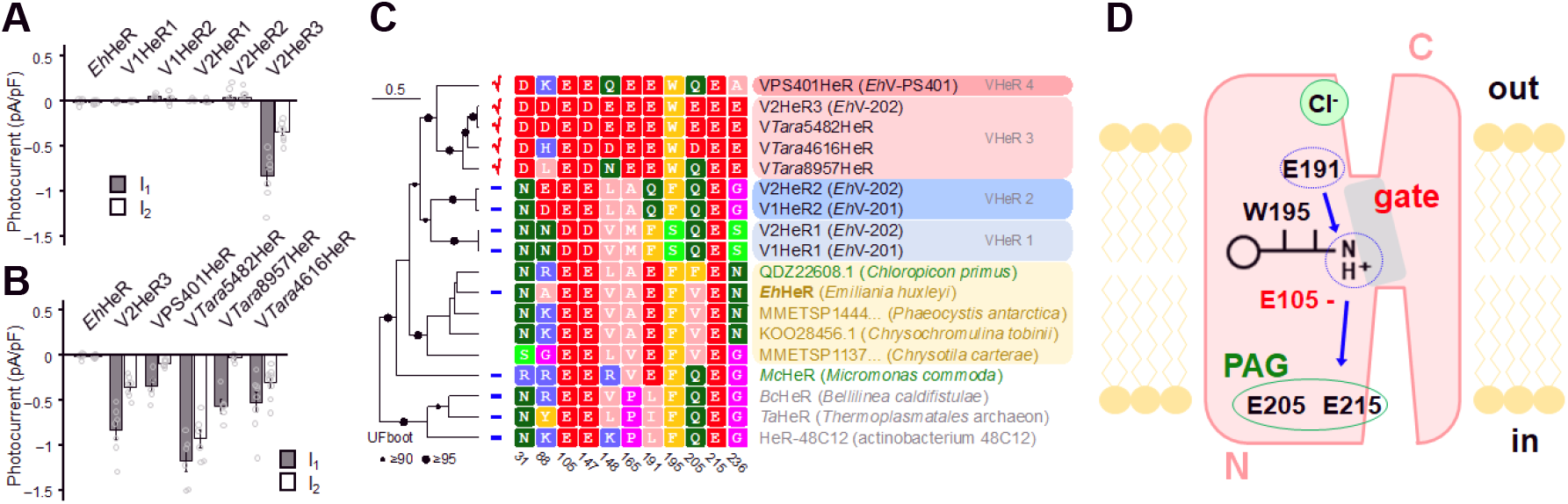
Electrophysiological measurements of related proteins of V2HeR3 and a functional model. (A) Electrophysiological measurements of a HeR from the host *E. huxleyi* (Ehux), two HeRs from *E. huxleyi* virus 201 (V1HeR1, 2) and three HeRs from *E. huxleyi* virus 202 (V2HeR1-3). Although some HeRs exhibit transient photocurrents (positive or negative peaks), steady-state photocurrent was only observed for V2HeR3. Comparison of photocurrent amplitudes at −40 mV. Square-block bar graph indicates amplitude from peak photocurrent, open bar graphs indicate amplitude from steady-state photocurrent. The pipette solution was 110 mM NaCl, pHi 7.4, the bath solution was 140 mM NaCl, pHe 7.4. n = 5 to 8 cells. (B) Electrophysiological measurements and the obtained I-V plots of homologous proteins of V2HeR3. Comparison of photocurrent amplitudes at −40 mV. Square-block bar graph indicates amplitude from peak photocurrent, open bar graphs indicate amplitude from steady-state photocurrent. The pipette solution was 110 mM NaCl, pHi 7.4, the bath solution was 140 mM NaCl, pHe 7.4. n = 5 to 8 cells. (C) Key residues for ion-transport of HeRs. (D) Schematic drawing of suggested proton-transporting mechanism in V2HeR3.

## Discussion

Fig. 4D outlines the proposed mechanism of proton channelling and pumping among the viral HeRs. Inward proton pump was recently found in nature (Harris et al., 2020; Inoue et al., 2020, 2016; Shevchenko et al., 2017), but the mechanism should be compared with outward proton pumps like BR, as the membrane topology is opposite to that of type-1 and -2 rhodopsins (Fig. 1B). Crystal structures of *Ta*HeR and 48C12 showed water-containing hydrogen-bonding networks between the retinal Schiff base and cytoplasmic aqueous phase. While the interior of the extracellular half is entirely hydrophobic in *Ta*HeR (Shihoya et al., 2019) and 48C12 (Kovalev et al., 2020; Lu et al., 2020), the proton-transporting HeRs reported here contain polar residues in that region. In particular, E191 appears to be the key residue for the function, presumably constituting the channel gate together with W195. Although E191 is a prerequisite for ion transport, it does not mean that this residue is negatively charged. Rather, E191 is protonated in the unphotolyzed state, and its deprotonation occurs upon opening of the gate. This residue is analogous to D96 in TM3 of BR and E90 in TM2 of ChR2, both of which are protonated in the dark state with the transient deprotonation necessary for light-driven proton pumping and light-gated cation channelling activities, respectively (Braiman et al., 1988; Gerwert et al., 1989; Kuhne et al., 2019; Radu et al., 2009). Similarly, in the cation channelrhodopsin 2 from *Guillardia theta* (GtCCR2) that shares with BR the DTD motif in TM3, channel opening requires deprotonation of the D96 homolog (Sineshchekov et al., 2017). Mutation of E205 or E215 in V2HeR3 led to conversion of proton pumping direction which suggests that E205 and E215 constitute the proton accepting group (PAG) upon M formation of HeR. These residues correspond to Q213 and E227 in *Ta*HeR, and Q216 and E230 in 48C12, which constitute a water-containing hydrogen-bonding network (Kovalev et al., 2020; Lu et al., 2020; Shihoya et al., 2019). As HeRs contain a single counterion of the Schiff base, it cannot be the proton acceptor upon deprotonation of the Schiff base. Instead, in the case of V2HeR3, the E205/E215 region appears to serve as PAG that receives the proton from the Schiff base. Interestingly, when the PAG mechanism is disrupted by mutation, this protein is converted to an outward proton pump. We found that monovalent anions affect E_rev_ only at the extracellular side (Fig. S4), suggesting its binding to V2HeR3 (Fig. 4D).

The discovery of proton-transporting HeRs provides various insights regarding evolution, physiology, mechanism of action and application of this rhodopsin family. Phylogenetic analysis strongly indicates that *coccolithoviruses* acquired their HeR genes from their past algal hosts (see Fig. S16). The split into proton-transporting and non-transporting types took place early in the evolution of the viral HeRs, while their distribution among *coccolithoviruses* suggests that the last common ancestor of all known isolates might have already possessed genes of both types (Fig. S17). The proton-transporting HeR was then secondarily lost in the lineage of *Eh*V-201. Proton-transporting HeRs is thus an innovation that appeared after the viruses acquired HeRs from algae, but the exact benefit that this function provided to the viruses is unclear. It was reported that light is required for viral adsorption to *E. huxleyi* cells (Thamatrakoln et al., 2019). If V2HeR3 is expressed in *E. huxleyi* cell membranes, light might depolarize the host cell membranes by this protein. It can be speculated that depolarization helps the virus to overcome cell defense or prevents superinfection. As *E. huxleyi* blooms significantly affect the marine environment and climate, the influence of light is intriguing. The viral HeRs are the first rhodopsin transporters to be characterized from the family Phycodnaviridae, paralleled by the type-1 rhodopsin channels in the Mimiviridae (Rozenberg et al., 2020; Zabelskii et al., 2020).

Type-1 microbial rhodopsins’ functions are diverse, while we now know that some HeRs participate in ion-transport such as proton channel and pump. The functional conversion of V2HeR3 into an outward proton pump may suggest the presence of such HeR proteins in nature. Thus, HeR’s functions are likely to be diverse similarly to type-1 rhodopsins. Finally, type-1 ion-transporting rhodopsins have been used as the main tools in optogenetics (Boyden et al., 2005; Deisseroth and Hegemann, 2017; Ishizuka et al., 2006), and ion-transporting HeRs will be potential optogenetic tools as well.

## Methods

### Sequence extraction

HeR genes from *Eh*V-202, *Eh*V-201 and *Eh*V-PS401 were obtained from the Genebank genome assemblies HQ634145.1 (Nissimov et al., 2012a), JF974311.1 (Nissimov et al., 2012b) and HQ634146.1 (Bioproject PRJNA47633), respectively: AET42597.1 (V2HeR1), AET42570.1 (V2HeR2) and AET42421.1 (V2HeR3) for *Eh*V-202, AET97940.1 (V1HeR1), AET97964.1 (V1HeR2) for *Eh*V-201 and AET73409.1 (VPS401HeR) for *Eh*V-PS401. Sequences related to the viral HeRs were searched for in the metagenomic databases Ocean Microbial Reference Catalog v.1 and v.2 (Villar et al., 2018), as well as other assemblies of the Tara Oceans data (Philosof et al., 2017; Sunagawa et al., 2015), with blastp v. 2.11.0+ (Altschul et al., 1990). After removing partial and redundant sequences, three full-length genes with a varying degree of relatedness to V2HeR3 were retained coming from contigs SAMEA2621401_1124616, SAMEA2621075_258957 and TARA_B100000767_G_C19749265_1 (containing OM-RGC gene OM-RGC.v1.010885482). The corresponding genes were dubbed V*Tara*4616HeR, V*Tara*8957HeR and V*Tara*5482HeR, respectively. Viral origin of the contigs is supported by matches to *Eh*Vs (Table S2). Annotated versions of the contigs are provided in Supplementary Dataset 1. The sequence used to represent the HeR gene from the host species (*Eh*HeR) was obtained from transcriptome assemblies of *Emiliania huxleyi* str. PLY M219 (corresponding to NCBI TSA transcript HBOB01045747.1). The gene is putatively single-copy, but has several splice variants and is detected in transcriptome assemblies of several *E. huxleyi* strains with minor allelic variation (Table S3). The gene is absent from the genome assembly of *E. huxleyi* CCMP1516 (NCBI Assembly GCA_000372725.1). Pairwise protein sequence identities are provided in Table S1. The identities were obtained by extracting transmembrane regions predicted for the full set of HeRs from the viruses and algae (see below) with PolyPhobius (Käll and Krogh, 2005).

### Phylogenetic reconstructions

Sequences for the analysis of the relationships between the viral and the eukaryotic HeRs were collected by searching a collection of 1315 transcriptomes and genomes from algae and other unicellular eukaryotes from NCBI Assembly, NCBI TSA, MMETSP (Johnson et al., 2019; Keeling et al., 2014), 1KP (Leebens-Mack, 2019), reefgenomics.org (Liew et al., 2016), as well de novo assemblies of data from NCBI SRA. If not annotated in the source databases, genes were predicted using GeneMark-ES v. 4.62 (Lomsadze et al., 2005) in the genomes or using Transdecoder v. 5.5.0 (https://github.com/TransDecoder) in the transcriptomes. Heliorhodopsin sequences were retrieved by searching the resulting protein database with hmmsearch from HMMER v. 3.3.2 (Eddy, 1996) using the Pfam HeR profile PF18761.4 with an E-value threshold of 1e-5, resulting in 565 sequences. Proteins most similar to the viral HeRs were obtained by searching among the HeRs using blastp with viral HeRs as queries with an E-value threshold of 1e-10. The resulting 268 sequences were clustered at 90% protein identity using cdhit v. 4.8.1 (Li and Godzik, 2006), truncated and mis-annotated sequences were removed and the resulting representative sequences together with the viral HeRs were aligned with mafft v. 7.475 (--localpair --maxiterate 1000) (Katoh et al., 2002), trimmed with trimal v. 1.4.rev15 (-gt 0.9) (Capella-Gutiérrez et al., 2009) and the phylogeny was reconstructed with iqtree v. 2.1.2 (Minh et al., 2020). The tree was midpoint-rooted. For a comparison to the more distant HeRs from prokaryotes and from Micromonas commoda a selected set of *Eh*V sequences and related algal HeRs were taken for a phylogenetic reconstruction using the same strategy.

Phylogenetic relationships between Coccolithovirus isolates were reconstructed by collecting orthologous genes with proteinortho v. 6.0.25 (Lechner et al., 2011) with blastp as the search engine. Viruses Ectocarpus siliculosus Virus-1 and Feldmannia species virus 158 from the sister genus Phaeovirus were recruited as outgroups. Strictly single-copy orthogroups present in at least 14 of the 16 genomes were selected and protein sequences were aligned with mafft (--localpair --maxiterate 1000), trimmed with trimal (-automated1) and the phylogeny was reconstructed with iqtree2 with partitions. Details on the HeR genes and phylogenetic markers in the used genomes are available in Dataset S2. The history of HeR gene duplications and losses among the three Coccolithovirus lineages was reconstructed with Notung v. 2.9.1.5 (Chen et al., 2000) using default costs, under the assumption of no gene transfer between lineages.

An unrooted phylogenetic tree in Fig. 1A was constructed with MEGAX software. The protein sequences were aligned using MUSCLE (Edgar, 2004). The evolutionary history was inferred using the Neighbor-Joining method (Saitou and Nei, 1987) with bootstrap values based on 1,000 replications. Sequence data of the other rhodopsins were from the GenBank database.

### Expression plasmids for mammalian cells

The expression plasmid for V2HeR3 with epitope tags (cMyc) and/or eGFP. peGFP-P2A-V2HeR3 was created by the following procedure. A full-length, DNA sequence encoding V2HeR3 was purchased from GenScript Japan (Tokyo, Japan). The gene encoding V2HeR3 and peGFP-P2A or pCaMKIIa-eGFP-P2A vector were amplified by PCR reactions, and V2HeR1 was subcloned into a peGFP-P2A vector or pCaMKIIa-eGFP-P2A vector using an In-Fusion HD cloning kit (Takara Bio, Inc., Shiga, Japan) according to the manufacturer’s instructions. For the immunostaining experiment, N-QKLISEEDL-C (10 amino acids, cMyc epitope tag) in the N-terminal or C-terminal of V2HeR3 was inserted in the plasmid peGFP-P2A-V2HeR3 using inverse PCR (KOD-Plus-Mutagenesis Kit, TOYOBO, Osaka, Japan). Site-directed mutagenesis was performed using a KOD-Plus-Mutagenesis Kit according to the manufacturer’s instructions.

Synthesized genes encoding *Eh*HeR, V1HeR2, V2HeR2, VPS401HeR, V*Tara*5482HeR, V*Tara*4616HeR and V*Tara*8957HeR were subcloned into a peGFP-P2A vector using an In-Fusion HD cloning kit. V1HeR2 and V2HeR2 were purchased from GenScript. *Eh*HeR, VPS401HeR, V*Tara*5482HeR, V*Tara*4616HeR and V*Tara*8957HeR were purchased from GENEWIZ Japan (Azenta Life Sciences, Tokyo, Japan). V1HeR1 and V2HeR1 were synthesized by GENEWIZ and subcloned into peGFP-P2A vector by GENEWIZ.

All the constructs were verified by DNA sequencing (Fasmac Co., Ltd. Kanagawa, Japan). All the PCR primers used in this study were summarized in Table S5-6.

### Mammalian cell culture

The electrophysiological and cytochemistry assays of HeRs were performed on ND7/23 cells, hybrid cell lines derived from neonatal rat dorsal root ganglia neurons fused with mouse neuroblastoma (Wood et al., 1990). ND7/23 cells were grown on a collagen-coated coverslip in Dulbecco’s modified Eagle’s medium (FUJIFILM Wako Pure Chemical Corporation, Osaka, Japan) supplemented with 2.5 μM all-*trans* retinal, 5% fetal bovine serum under a 5% CO_2_ atmosphere at 37 °C. The expression plasmids were transiently transfected by using Lipofectamine 3000 (Thermo Fisher Scientific, Waltham, MA, USA) according to the manufacturer’s instructions. Electrophysiological recordings were then conducted 16-36 h after the transfection. Successfully transfected cells were identified by eGFP fluorescence under a microscope prior to the measurements.

Cortical neurons were isolated from embryonic day 16 Wistar rats (Japan SLC, Inc., Shizuoka, Japan) using Nerve-Cells Dispersion Solutions (FUJIFILM Wako Pure Chemical Corporation) according to the manufacturer’s instructions and grown in culture medium (FUJIFILM Wako Pure Chemical Corporation) under a 5% CO_2_ atmosphere at 37 °C. The expression plasmids were transiently transfected in cortical neurons calcium phosphate transfection at days in vitro (DIV) 5. Electrophysiological recordings were then conducted at DIV21-23 to neurons identified to express eGFP fluorescence under conventional epifluorescence system.

### Electrophysiology

All experiments were carried out at room temperature (23±2°C). Photocurrents and action potentials were recorded as previously described using an Axopatch 200B amplifier (Molecular Devices, Sunnyvale, CA, USA) under a whole-cell patch clamp configuration (Hososhima et al., 2021). The data were filtered at 5 kHz and sampled at 20 kHz (Digdata1550, Molecular Devices, Sunnyvale, CA, USA) and stored in a computer (pClamp10.6, Molecular Devices). The pipette resistance was between 3–10 MΩ. All patch-clamp solutions are described in Table S7. The liquid junction potential was calculated and compensated by the pClamp 10.6 software.

For whole-cell patch clamp, irradiation at 470 or 530 nm was carried out using WheeLED (parts No. WLS-LED-0470-03 or WLS-LED-0530-03, Mightex, Toronto, Canada) controlled by computer software (pCLAMP10.6, Molecular Devices). The light power was directly measured at an objective lens of microscopy by a visible light-sensing thermopile (MIR-100Q, SSC Inc., Mie, Japan).

Transient photocurrent (I_0_) corresponds to initial peak photocurrents during light illumination, peak photocurrent (I_1_) corresponds to average value of photocurrents from 10-11 ms, steady-state photocurrent (I_2_) corresponds to average value of photocurrents of the pulse-end 10 ms. All data in the text and figures are expressed as mean ± SEM and were evaluated with the Mann-Whitney *U* test for statistical significance, unless otherwise noted. It was judged as statistically insignificant when P > 0.05.

### Cytochemistry

The cultured ND7/23 cells on glass coverslips were washed with PBS (NACALAI TESQUE, INC., Kyoto, Japan). Two cell samples were fixed in 4% paraformaldehyde phosphate buffer solution (NACALAI TESQUE, INC.) for 15 min at room temperature. The cells were washed with PBS three times. One sample was permeabilized with 0.5% Triton X-100 for 15 min at room temperature, the other sample was not permeabilized. The cells were treated with blocking buffer consisting of 3% goat serum for 60 min at room temperature. Then the cells were reacted with rabbit anti–c-Myc primary antibody (C3956; Sigma-Aldrich, St. Louis, MO, USA) at 1:500 dilution for 60 min at room temperature. The cells were washed with PBS three times before labeling with goat anti-rabbit IgG H&L Biotin (ab97049; abcam, Cambridge, UK) at 1:500 dilution for 30 min at room temperature. After that the cells were washed PBS three times before labeling with streptavidin, Alexa Fluor 594 (S32356; Thermo Fisher Scientific) at 1:200 dilution for 2 hours at room temperature. After a final wash with PBS three times, the coverslips were mounted on glass slides with ProLong Diamond Antifade Mountant (Thermo Fisher Scientific). Localization was assessed using a LSM880 confocal laser scanning microscope (Zeiss, Jena, Germany) equipped with ×63 oil-immersion objective lens (Zeiss) and a software ZEN (Zeiss). The captured images were analyzed with Fiji software (Schindelin et al., 2012).

### Expression plasmids for Pichia pastoris cells

For the purification of V2HeR3-WT, the gene encoding N-terminal His-tagged V2HeR3 was cloned into the EcoRI and XbaI site of pPICZB vector (Thermo Fisher Scientific). For determination of λmax of V2HeR3 WT and mutants, the S-tag (KETAAAKFERQHMDS) and thrombin recognized sequence (LVPRGS) were added between N-terminal His-tag and V2HeR3 coding region.

### Protein expression and purification by Pichia pastoris cells

The gene encoding N-terminal His-tagged V2HeR3 was cloned into the EcoRI and XbaI site of pPICZB vector (Thermo Fisher Scientific). The recombinant protein was expressed in the *Pichia pastoris* strain SMD1168H (Philosof et al., 2017) (Thermo Fisher Scientific). The cells were harvested 48–60 h after expression was induced in BMMY medium when 10 mM of all-*trans*-retinal (Sigma-Aldrich) was supplemented in the culture to a final concentration of 30 μM. Additionally, 100% filtered methanol was added to the growth medium every 24 h of induction to a final concentration of 0.5 %. Membranes containing V2HeR3 was isolated as described elsewhere (Yamauchi et al., 2017) with the following modifications. Washed *P. pastoris* cells were resuspended in buffer A (7 mM NaH_2_PO_4_, 7 mM EDTA, 7 mM DTT, and 1 mM phenylmethylsulfonyl fluoride (PMSF), pH 6.5) and slowly shaken with all-*trans*-retinal (added to a final concentration of 25 μM) in the dark at room temperature for 3–4 h in the presence of 0.5 % of Westase (Takara Bio, Inc.) to digest the cell wall. The cells were disrupted by the 2-times passage through a high-pressure homogenizer (EmulsiFlex C3, Avestin, Inc., Canada). The supernatants were centrifuged for 30 min at 40,000×g in a fixed-angle rotor, and the V2HeR3 membrane pellets were resuspended in solubilization buffer (20 mM KH_2_PO_4_, 1 % n-dodecyl-β-D-maltoside (DDM), 1 mM PMSF, pH 7.5) and stirred overnight at 4°C. The solubilization mixture was centrifuged for 30 min at 40,000×g in a fixed-angle rotor. The solubilized protein was incubated with Ni-NTA agarose (QIAGEN, Hilden, Germany) for several hours. The resin with bound V2HeR3 was washed with wash buffer (50 mM KH_2_PO_4_, 400 mM NaCl, 0.1 % DDM, 35 mM imidazole, pH 7.5) and then treated with elution buffer (50 mM KH_2_PO_4_, 400 mM NaCl, 0.1 % DDM, 250 mM imidazole, pH 7.5). The collected fractions were dialyzed against a solution containing 50 mM KH_2_PO_4_, 400 mM NaCl, 0.1 % DDM at pH7.5 to remove the imidazole.

### Ion transport assay of Pichia pastoris cells by pH electrode

The number of *P. pastirs* cells expressing rhodopsins was estimated by the apparent optical density at 660 nm (OD_660_), and 7.5 ml cell culture (OD_660_ = 2) were used for the experiment. The cells were washed with an unbuffered 100 mM NaCl solution three times, and resuspended in the same solution. The cell suspension was placed in the dark and then illuminated at λ > 500 nm, by the output of a 1-kW tungsten–halogen projector lamp (Rikagaku, Japan) through a glass filter (Y-52, AGC Techno Glass, Japan). The light-induced pH changes were measured with a pH electrode (HORIBA, Ltd, Japan) (Inoue et al., 2013). Measurements were repeated under the same conditions with the addition of 30 μM CCCP, a protonophore molecule.

### HPLC analysis of retinal configuration

The high-performance liquid chromatograph was equipped with a silica column (6.0 × 150 mm; YMC-Pack SIL, YMC, Japan), a pump (PU-2080, JASCO, Japan), and a UV-visible detector (UV-2070, JASCO) (Kawanabe et al., 2006). The solvent was composed of 12% (v/v) ethyl acetate and 0.12% (v/v) ethanol in hexane with a flow rate of 1.0 ml min^−1^. Retinal oxime was formed by a hydrolysis reaction with the sample in 100 μl solution at 0.1 mg ml^−1^ protein concentration and 50 µl hydroxylamine solution at 1 M at 0°C. To ensure all the protein molecules reacted completely, 300 μl of methanol was added to denature the proteins. For light-adapted V2HeR3, the sample solution was illuminated with λ > 500 nm light (Y-52, AGC Techno Glass) for 1 min before denaturation and extraction. Then, the retinal oxime was extracted using hexane and 300 μl of solution was injected into the HPLC system. The molar composition of the retinal isomers was calculated from the areas of the corresponding peaks in the HPLC patterns. The assignment of the peaks was performed by comparing them with the HPLC pattern from retinal oximes of authentic all-*trans*, 13-*cis*, and 11-*cis* retinals. To estimate the experimental error, three independent measurements were carried out.

### pH titration

To investigate the pH dependence of the absorption spectra of V2HeR3, a solution containing about 6 μM protein was solubilized in 6-mix buffer (10 mM citrate, 10 mM MES, 10 mM HEPES, 10 mM MOPS, 10 mM CHES, and 10 mM CAPS). The pH was then changed by the addition of concentrated HCl or NaOH (7). The absorption spectra were measured with a UV-visible spectrometer (V-2400PC, SHIMADZU, Japan) at each approximately 0.5-pH change.

### Laser flash photolysis

For the laser flash photolysis measurement, V2HeR3 was purified and solubilized in 0.1% DDM, 400 mM NaCl and 50 mM KH_2_PO_4_ (pH 7.5). The absorption of the protein solution was adjusted to 0.5 (total protein concentration ∼0.25 mg mL^−1^) at λ_max_ = 500 nm. The sample was illuminated with a beam from an OPO system (LT-2214, LOTIS TII, Minsk, Republic of Belarus) pumped by the third harmonics of a nanosecond pulsed Nd^3+^-YAG laser (λ = 355 nm, LS-2134UTF, LOTIS TII) (61). The time-evolution of transient absorption change was obtained by observing the intensity change of an output of an Xe arc lamp (L9289-01, Hamamatsu Photonics, Japan), monochromated by a monochrometer (S-10, SOMA OPTICS, Japan) and passed through the sample, after photo-excitation by a photomultiplier tube (R10699, Hamamatsu Photonics, Japan). To increase the signal-to-noise (S/N) ratio, multiple measurements were averaged. The signals were global-fitted with a multi-exponential function to obtain the lifetimes of each photo-intermediate.

### Low-temperature UV-visible and FTIR spectroscopy

The purified proteins of V2HeR3 were reconstituted into a mixture of POPE and POPG membranes (molar ratio = 3:1) with a protein-to-lipid molar ratio of 1:20 by removing DDM using Bio-Beads (SM-2; Bio-Rad, CA, USA). The reconstituted samples were washed three times with 1 mM NaCl and 2 mM Tris-HCl (pH 8.0). The pellet was resuspended in the same buffer, where the concentration was adjusted to make the intensity of amide I ∼ 0.7. A 60-μL aliquot was placed onto a BaF_2_ window and dried gently at 4°C. The films were then rehydrated with 2 μL H_2_O or D_2_O, and allowed to stand at room temperature for 15 min to complete the hydration. For UV-visible spectroscopy, the sample film was hydrated with H_2_O, and placed and cooled in an Optistat DN cryostat mounted (Oxford Instruments, Abingdon, UK) in a UV-vis spectrometer (V-550, JASCO, Japan) (Kawanabe et al., 2007). For FTIR spectroscopy, the sample film was hydrated with H_2_O or D_2_O, and placed and cooled in an Oxford Optistat DN2 cryostat mounted in a Cary670 spectrometer (Agilent Technologies, Japan) (Shihoya et al., 2019). The 128 interferograms were accumulated with 2 cm^−1^ spectral resolution for each measurement.

For the formation of the K intermediate, samples of V2HeR3 were illuminated with 520-nm light (interference filter) from a 1 kW tungsten–halogen projector lamp (Rikagaku) for 2 min at 100 K. The K intermediate was photo-reversed with λ > 590 nm light (R-61 cut-off filter, Toshiba, Japan) for 1 min, followed by illumination with 540-nm light. For the formation of the M intermediate, samples of V2HeR3 were illuminated with λ > 500 nm light (Y-52 cut-off filter, Toshiba) from a 1 kW halogen-tungsten lamp for 1 min at 230 K. The M intermediate was photo-reversed with the 400-nm light (interference filter) for 2 min, followed by illumination with λ > 500 nm light. To increase the signal/noise ratio in FTIR spectroscopy, photoconversions to the K intermediate at 100 K and to the M intermediate at 230 K were repeated 8 and 5 times, respectively.

### Determination of λ_max_ by hydroxylamine bleach

The λ_max_ values of HeRs in ND7/23 or *P. pastoris* cells were determined by observing the bleaching by hydroxylamine upon light absorption. ND7/23 cells were grown as mentioned above. The expression plasmids were transiently transfected by using PEI-Max according to the manufacturer’s instructions. ND7/23 cells 100 mm-dish x2) expressing rhodopsin was centrifuged and resuspended in 50 mM Tris-Cl (pH 8.0), 100 mM NaCl buffer to a final volume of 0.7 ml. Then, the mammalian cells were disrupted by ultrasonication and solubilized in 1.0% DDM. We added hydroxylamine to the sample (final concentration of 50 mM) and illuminated it for 16 min with a SOLIS-1C -High-Power LED (THORLABS, Japan) through a glass filter (Y-52, AGC Techno Glass) at wavelengths > 500 nm. Absorption changes representing the bleaching of rhodopsins by hydroxylamine were measured, using a UV-visible spectrometer, V750 (JASCO) with integrating sphere unit, ISV922.

*P. pastoris* membranes expressing rhodopsin was resuspended in 50 mM Tris-Cl (pH 8.0), 100 mM NaCl buffer to final concentration 5 mg/ml. Then, membrane fraction solubilized in 1.0% DDM. We added hydroxylamine to the sample (final concentration of 500 mM) and illuminated it for 16 min with a 1 kW tungsten-halogen projector lamp (Master HILUX-HR, Rikagaku) through a glass filter (Y-52, AGC Techno Glass) at wavelengths > 500 nm. Absorption changes representing the bleaching of rhodopsins by hydroxylamine were measured, using a UV-visible spectrometer V650 (JASCO) with integrating sphere unit.

## Acknowledgments

This work was financially supported by grants from the Japanese Ministry of Education, Culture, Sports, Science and Technology (18H03986, 19H04959, 20K21251, 21H04969 to H.K.; 18K06109 to S.P.T.; 17H03007 to K.I.; 20K15900 to S.H.); and Israel Science Foundation (F.I.R.S.T. program 3592/19 and Research Center grant 3131/20 to O.B.), O.B. holds the Louis and Lyra Richmond Chair in Life Sciences.

## Author Contributions

A.P., O.B. and H.K. conceived the project, and H.K. coordinated the project. S.H., R.M., R.A.Y., and M.K. performed molecular biology and protein biochemistry; S.H., S.S., and performed electrophysiology; R.M., K.M., and K.I. performed laser flash photolysis and HPLC analysis; R.M. and K.K. performed FTIR spectroscopy; S.H. and R.A.Y. performed hydroxylamine bleach; A.R., and O.B. performed bioinformatic analyses; H.K. wrote the paper, which was critically revised and approved by all authors.

## Supplementary Information

### Supplementary Information section

**Fig. S1.**
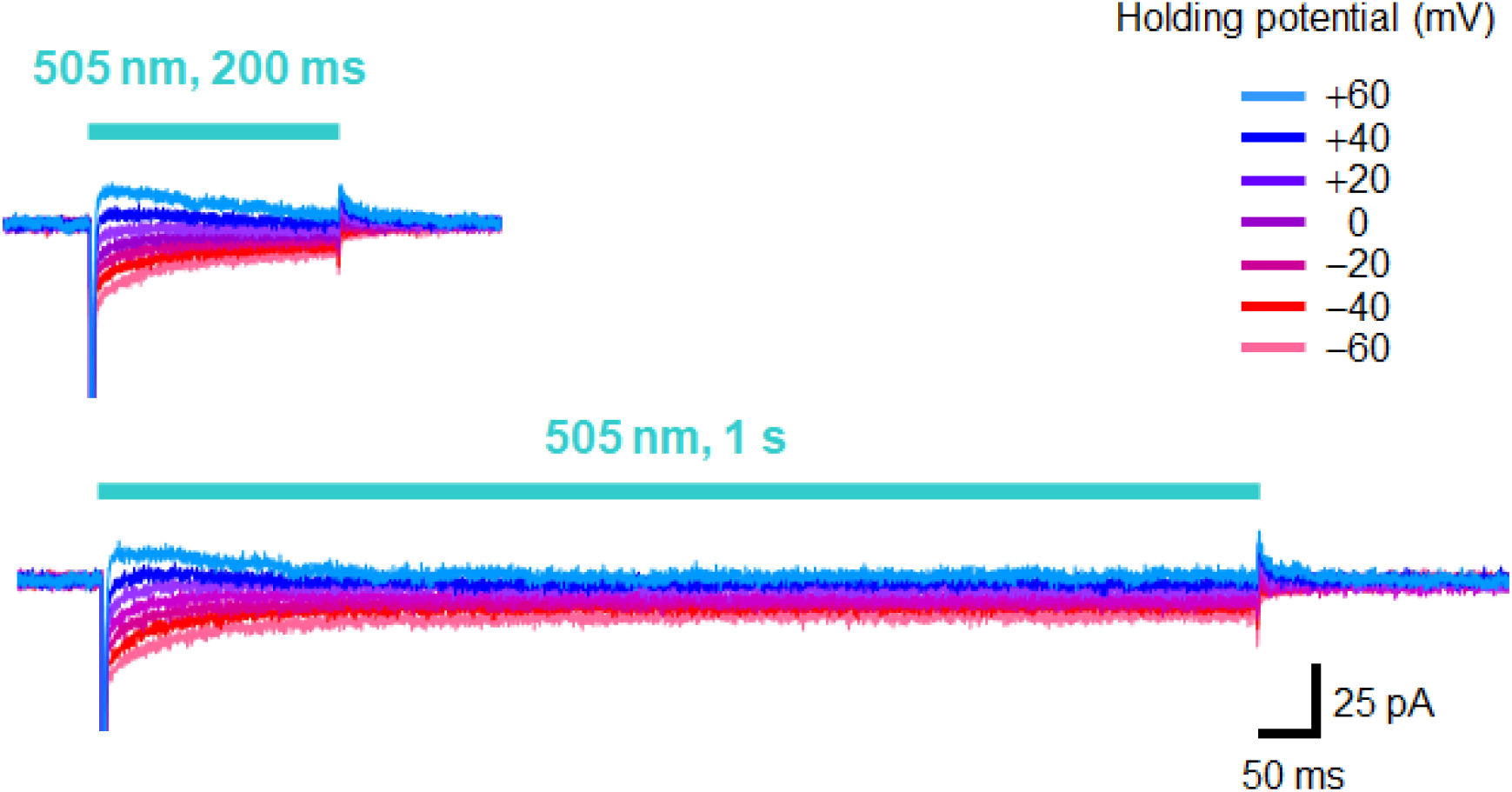
Photocurrents of V2HeR3 under different illumination periods. Electrophysiological measurements of V2HeR3-driven photocurrent in ND7/23 cells. The cells were illuminated with light (λ = 505 nm, 24.5 mW/mm^2^) during the time region shown by blue bars. The membrane voltage was clamped from −60 to +60 mV for every 20-mV step. The pipette solution was 110 mM NaCl, pHi 7.4, the bath solution was 140 mM NaCl, pHe 7.4.

**Fig. S2.**
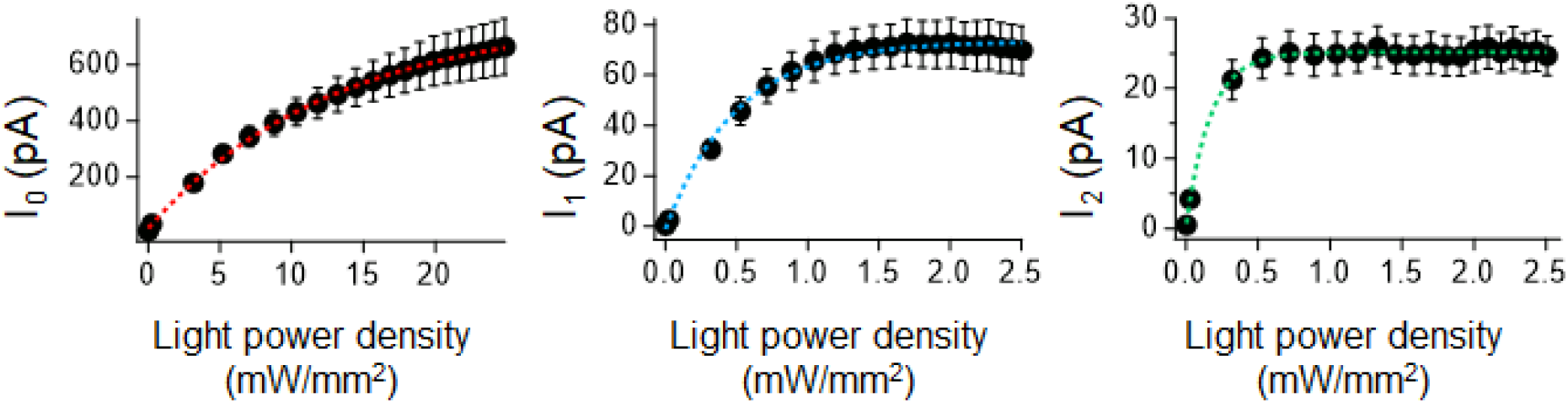
Photocurrents of V2HeR3 under different light intensities. Photocurrent amplitude was plotted as a function of light power. Membrane voltage was clamped at −60 mV, 505 nm light were illuminated respectively. Half saturation maxima (EC_50_) of the transient photocurrent (I_0_; left), peak photocurrent (I_1_; middle) and steady-state photocurrent (I_2_; right) are 13 ± 2.2 mW/mm^2^ (± SEM), 0.48 ± 0.017 mW/mm^2^ and 0.18 ± 0.016 mW/mm^2^, respectively. The pipette solution was 110 mM NMG-Cl, pHi 7.4, the bath solution was 140 mM NaCl, pHe 7.4. n = 5 to 8 cells.

**Fig. S3.**
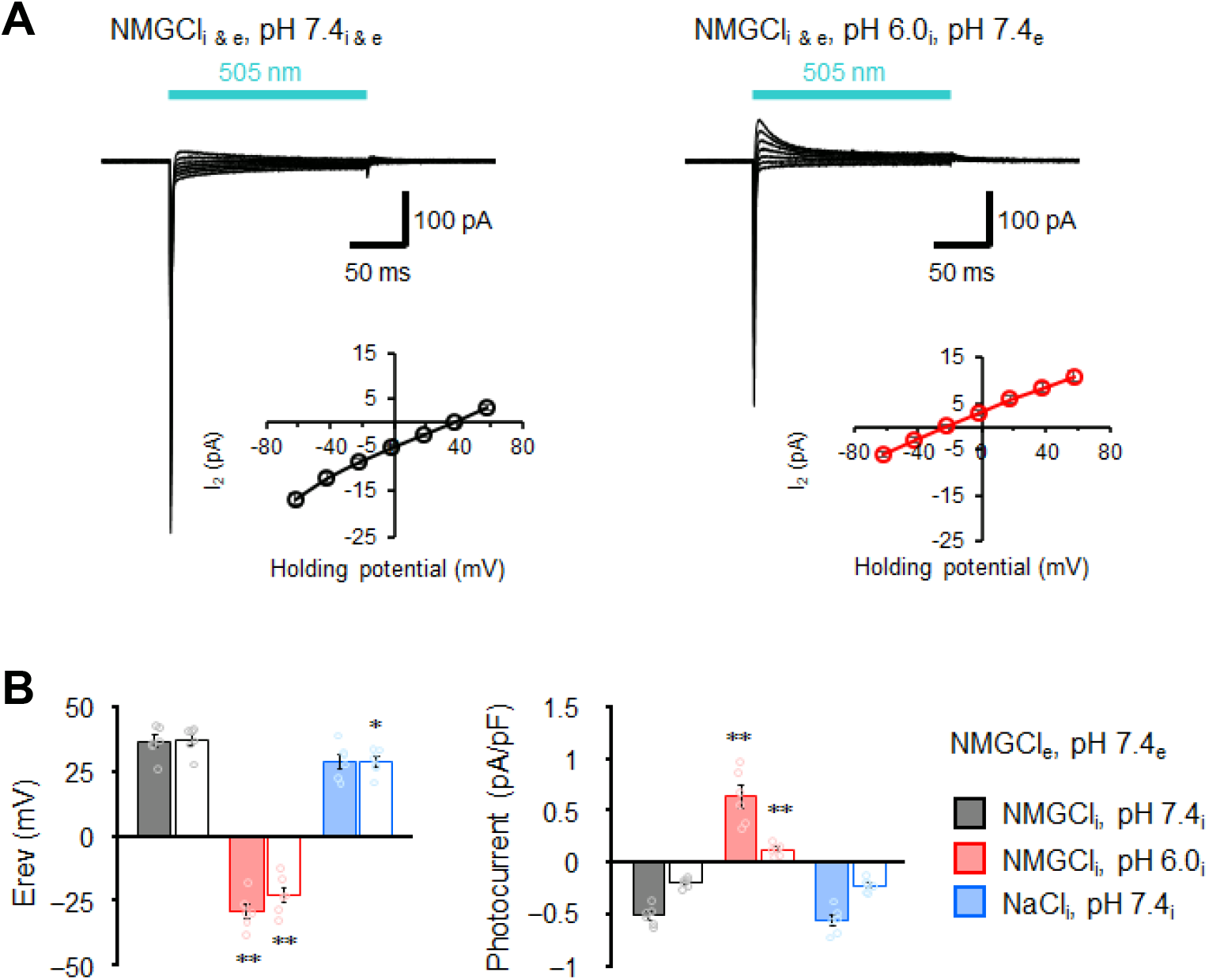
Selectivity of cations in the transport of V2HeR3. (A) Representative photocurrent trace of V2HeR3 recorded in ND7/23 cells. The cells were illuminated with light (λ = 505 nm, 24.5 mW/mm^2^) during the time region shown by blue bars. Voltage was clamped from −60 mV to +60 mV in 20-mV steps. The bath solution was 140 mM NMG-Cl, pHe 7.4. The pipette solution was 110 mM NMG-Cl, pHi 7.4 (left). The pipette solution was 110 mM NMG-Cl, pHi 6.0 (right). n = 6 cells. (B) Corresponding reversal potential (Erev) for each internal condition (left), and comparison of photocurrent amplitudes at 0 mV for different internal cations (right). Square-block bar graph indicates Erev or amplitude from peak photocurrent, open bar graphs indicate Erev or amplitude from steady-state photocurrent. n = 6 cells. (*p<0.05, ** p<0.01)

**Fig. S4.**
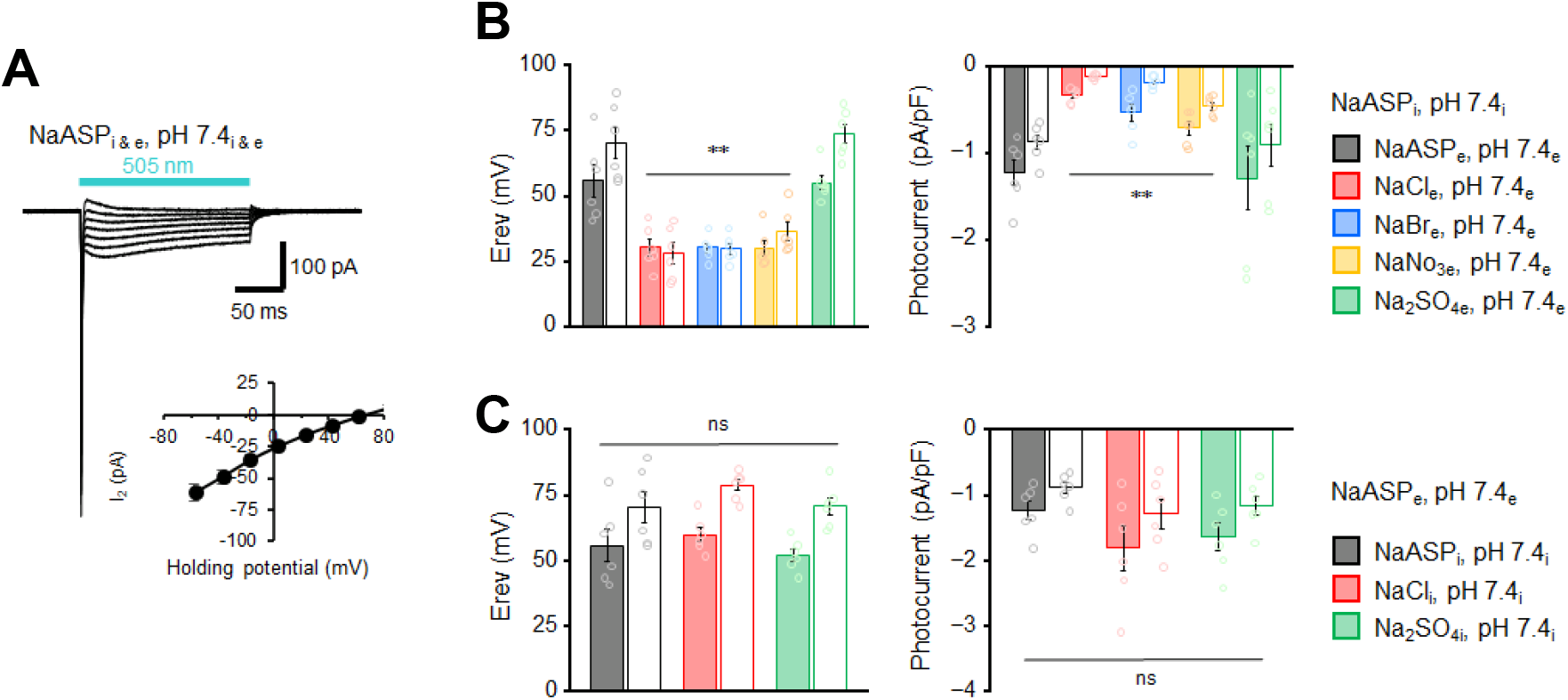
Selectivity of anions in the transport of V2HeR3. (A) Representative photocurrent trace of V2HeR3 recorded in ND7/23 cells. The cells were illuminated with light (λ = 505 nm, 24.5 mW/mm^2^) during the time region shown by blue bars. Voltage was clamped from −60 mV to +60 mV in 20-mV steps. The pipette solution was 110 mM NaASP, pHi 7.4, the bath solution was 140 mM NaASP, pHe 7.4. n = 6 cells. (B), (C) Corresponding reversal potential (E_rev_) for each external (B) or internal (C) condition (left), and comparison of photocurrent amplitudes at 0 mV for different external (B) or internal (C) anions (right). Square-block bar graph indicates E_rev_ or amplitude from peak photocurrent, open bar graphs indicate Erev or amplitude from steady-state photocurrent. n = 6 cells. (** p<0.01)

**Fig. S5.**
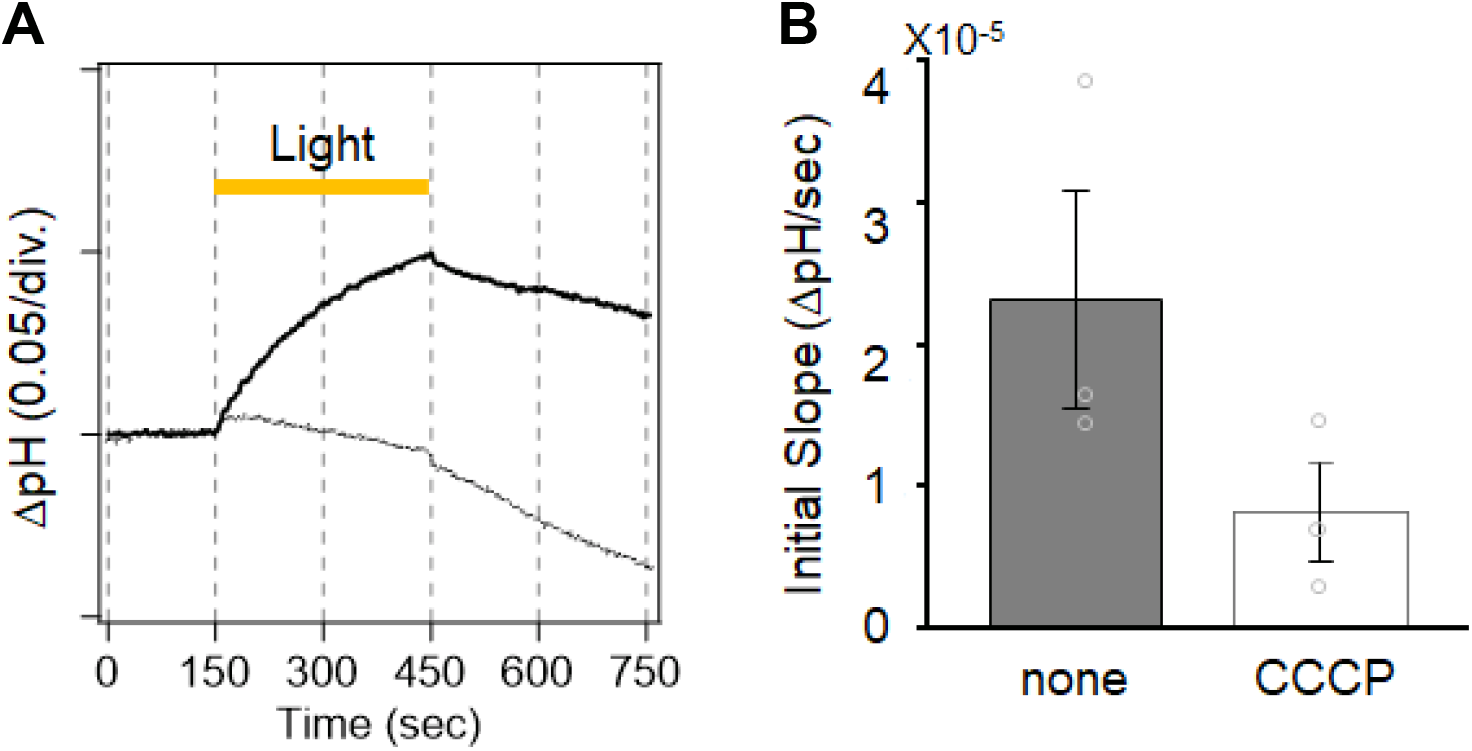
Ion transport activity of V2HeR3 in *Pichia pastirs*. (A) Pump activity of *P. pastris* cell suspensions (100 mM NaCl) expressing V2HeR3-WT without (solid line) and with (dotted line) CCCP. (B) Initial slope in light-induced pH changes.

**Fig. S6.**
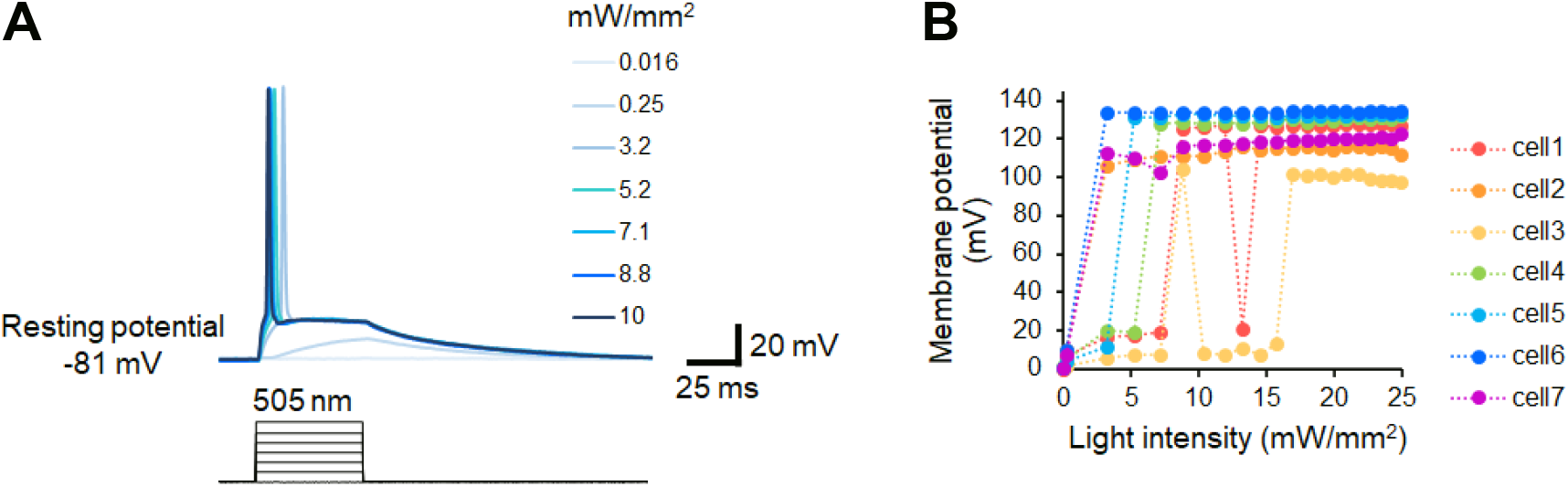
Light dependency of neuronal manipulation by V2HeR3. (A) Representative action potentials evoked by various light power. 505 nm light was applied in neurons for 50 ms as indicated by black bars. (B) Evoked action potentials from 7 neurons were plotted as a function of light power.

**Fig. S7.**
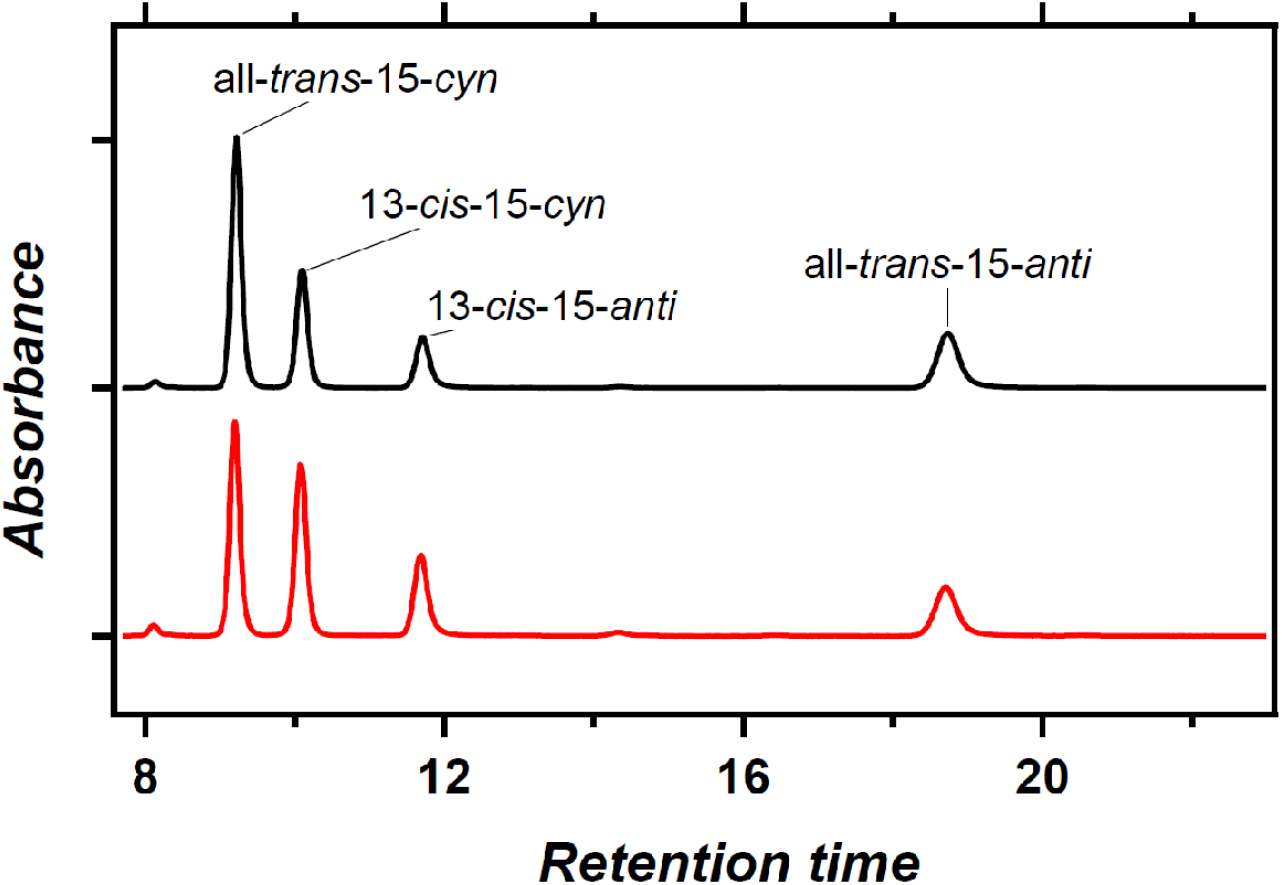
HPLC pattern of the retinal chromophore in V2HeR3. HPLC pattern of retinal extracted from V2HeR3 in the dark (black) and during light illumination at λ = 500 ± 10 nm (red). The populations of the all-*trans* and 13-*cis* forms are 64 % and 36 % in the dark, and 51 % and 49 % in the light, respectively.

**Fig. S8.**
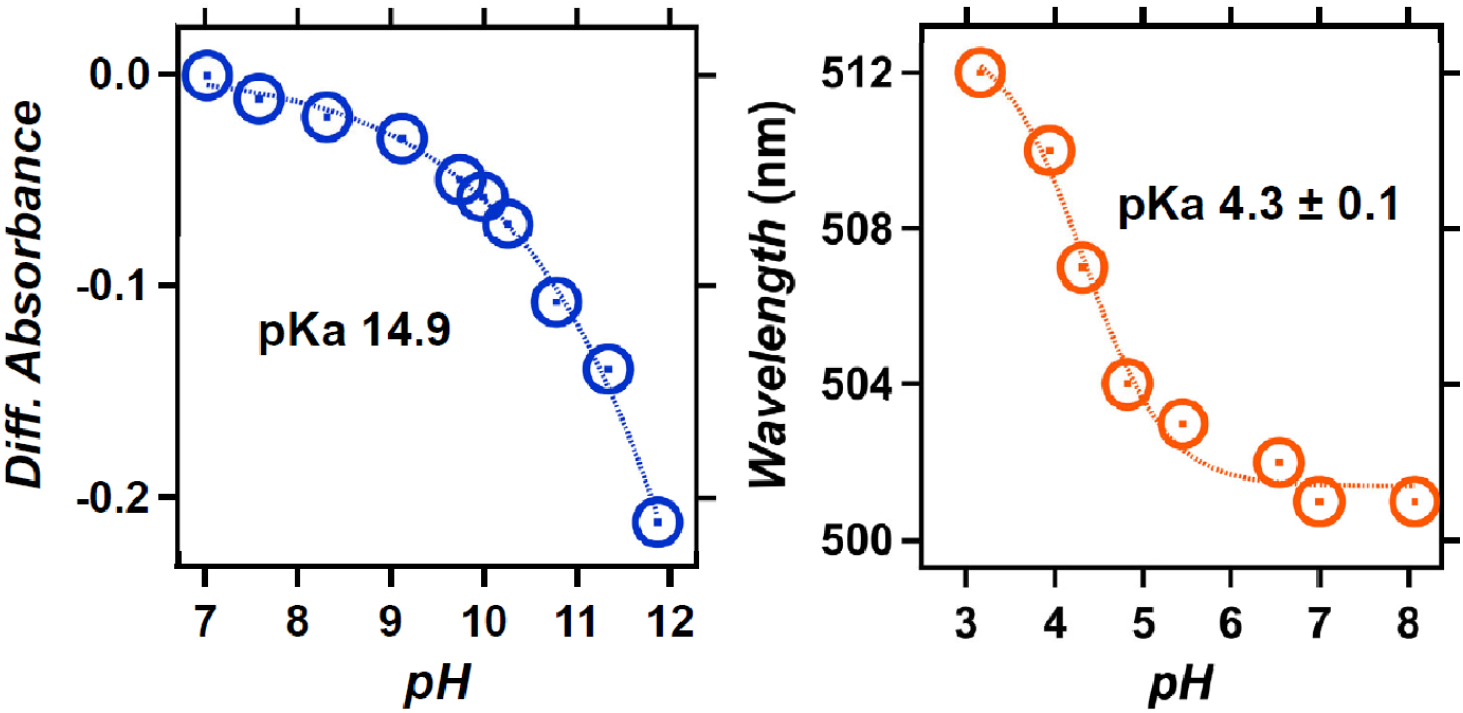
pH titration of V2HeR3. UV-visible absorption spectra (left) and the λ_max_ (right, orange solid circles) of heliorhodopsin 48C12 at pH 2.8-8.4. When pH is lowered, a red-shift of the absorption is observed, which is commonly reported for many type-1 rhodopsins and reflects protonation of counterions. Thus, the red-shift of heliorhodopsin 48C12 originates from protonation of E107, which is fitted with the Henderson–Hasselbalch equation (blue dashed line), and the pKa of counterion (E107) is estimated to be 3.7. At pH < 2.8, a large blue shift to 443 nm is observed, presumably owing to the acid denaturation of the protein.

**Fig. S9.**
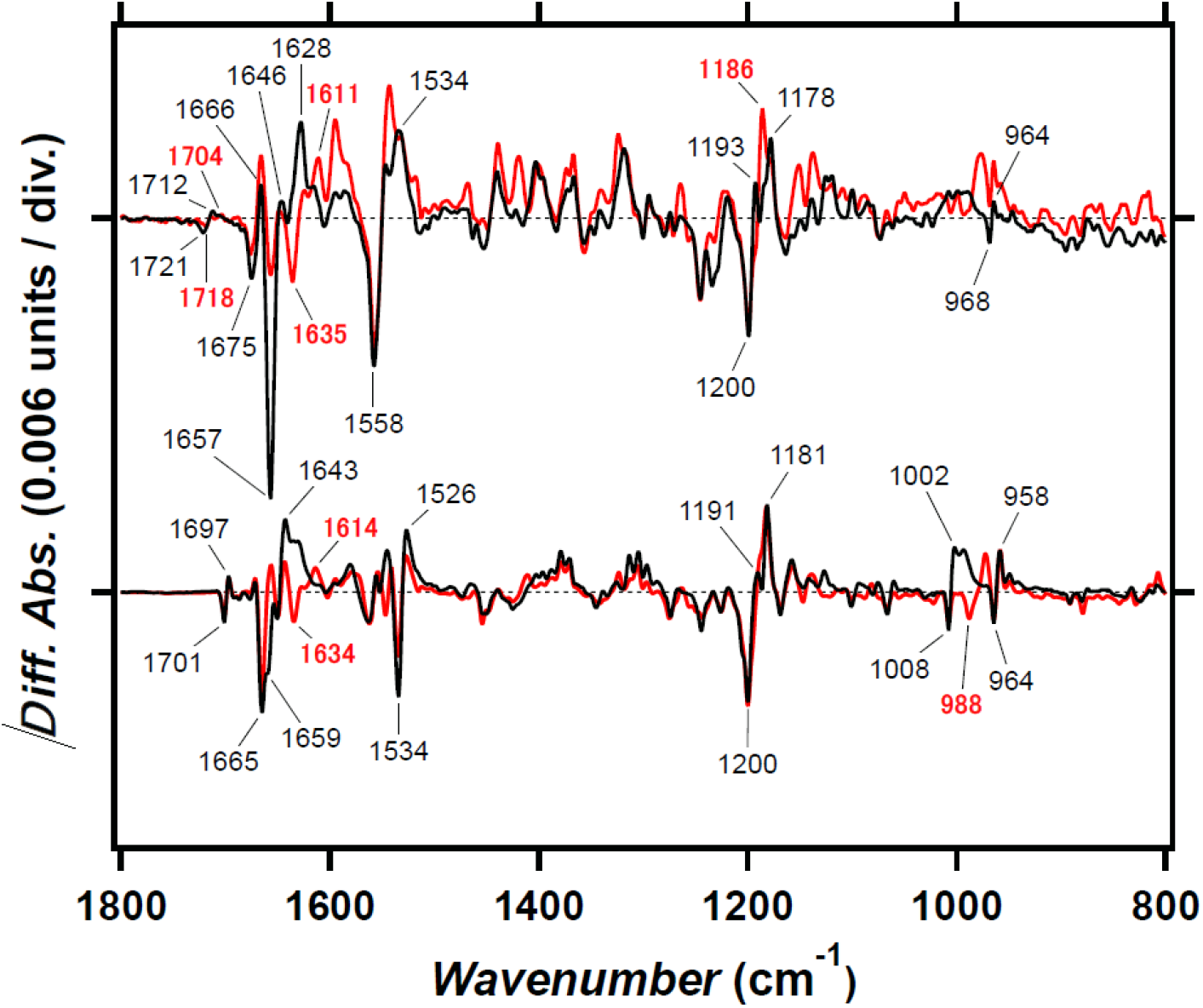
Light-induced difference FTIR spectra of V2HeR3 and *Ta*HeR at 77 K. Light-induced difference FTIR spectra of V2HeR3 (upper) and *Ta*HeR (lower) in the 1800-800 cm^-1^ region at 100 and 77 K, respectively, latter of which is reproduced from Ref.7. Positive and negative bands represent vibrations of the primary K intermediate and the unphotolyzed state, respectively, and characteristic vibrational bands are tagged. C=C stretch, C-C stretch, and hydrogen-out-of-plane (HOOP) vibrations of the retinal chromophore appear at 1550-1500, 1250-1100, and 1000-900 cm^-1^, respectively. These signals resemble between V2HeR3 and *Ta*HeR, suggesting common structural changes such as all-*trans* to 13-*cis* isomerization with distorted chromophore. Amide-I vibration of peptide backbone and carboxylic C=O stretch appear at 1700-1600, and 1800-1700 cm^-1^, respectively. Intense band at 1657 (−)/1628 (+) cm^-1^ in VeHeR3 is a signature of larger changes in the secondary structure in VeHeR3 than that in *Ta*HeR. Carboxylic C=O stretch vibrations appear only for V2HeR3 at 1721 (−)/1712 (+) cm^-1^, not for *Ta*HeR.

**Fig. Figure S10.**
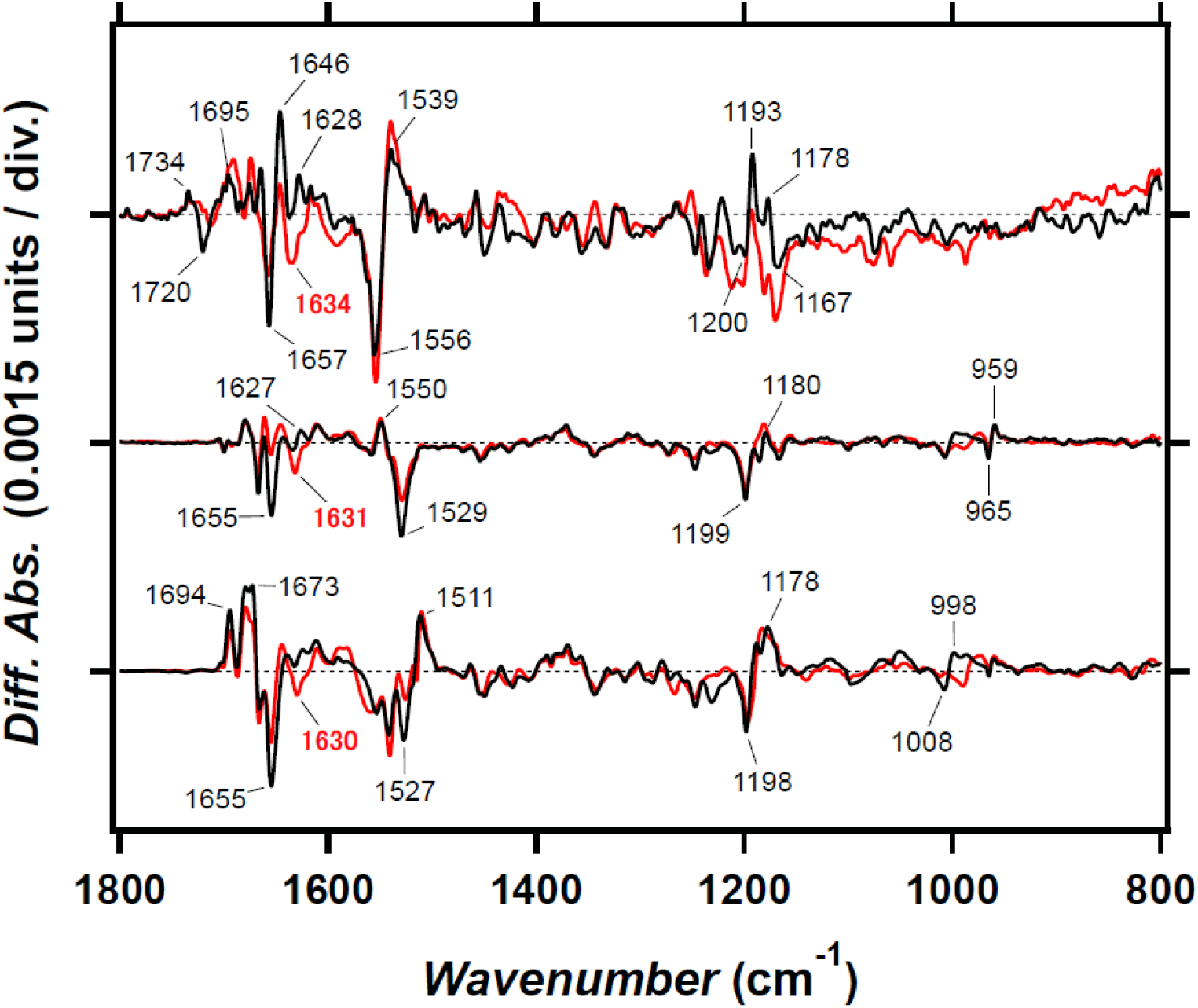
Light-induced difference FTIR spectra for late intermediates of V2HeR3 and *Ta*HeR. Light-induced FTIR spectra of V2HeR3 at 230 K (upper), *Ta*HeR at 240 K (middle), and *Ta*HeR at 277 K (lower) in the 1800-800 cm^-1^ region. The spectra of *Ta*HeR are reproduced from Ref.7. While negative bands correspond to vibrations of the unphotolyzed state, positive bands of the upper, middle, and lower panels correspond to the M intermediate of V2HeR3, the M intermediate of *Ta*HeR, and the O intermediate of *Ta*HeR, respectively. Strong amide-I band, observed at 1657 (−)/1646 (+) cm^-1^ in V2HeR3 (upper), is similar to the case in the O intermediate of *Ta*HeR (lower), not the M intermediate of *Ta*HeR (middle).

**Fig. Figure S11.**
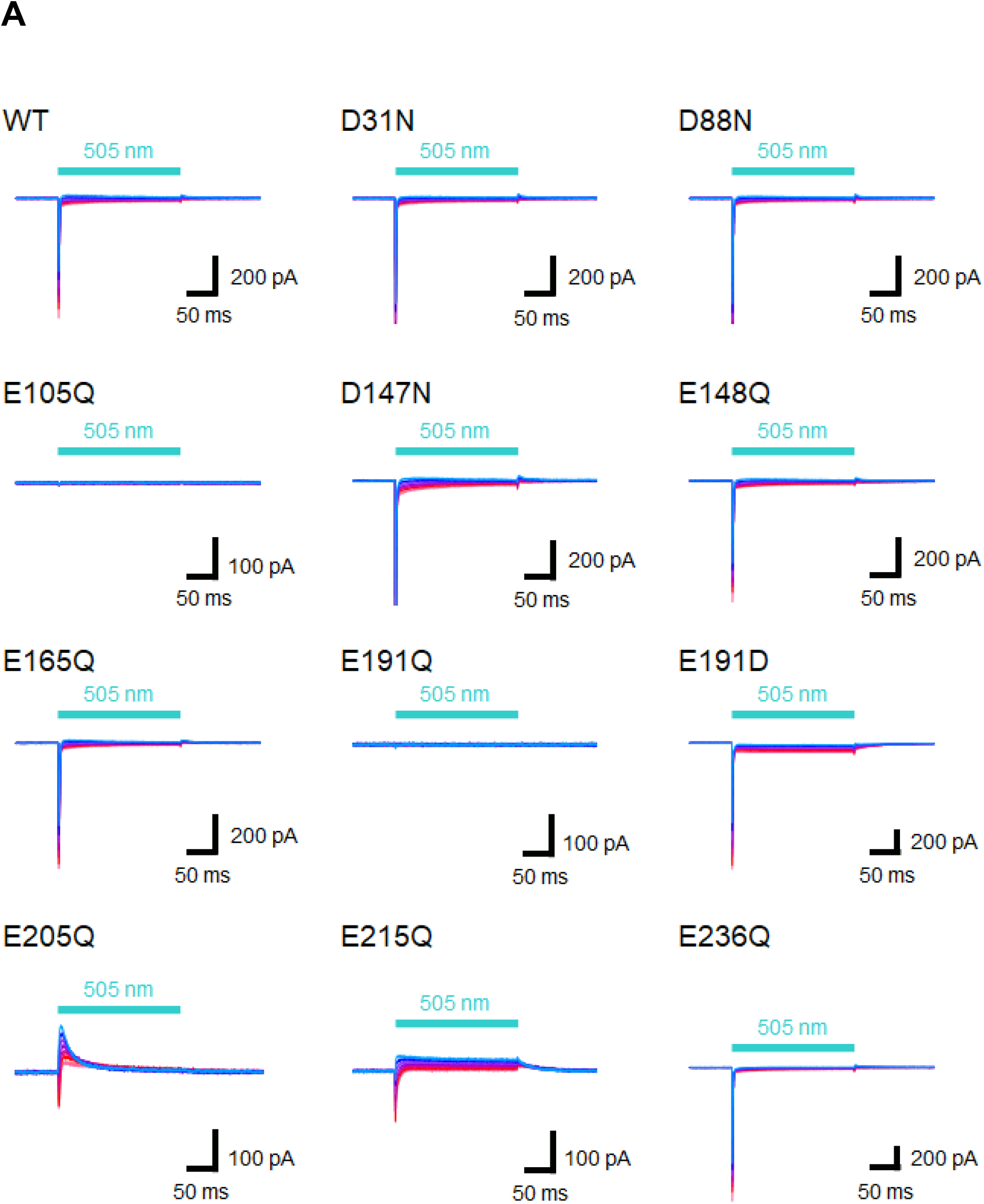
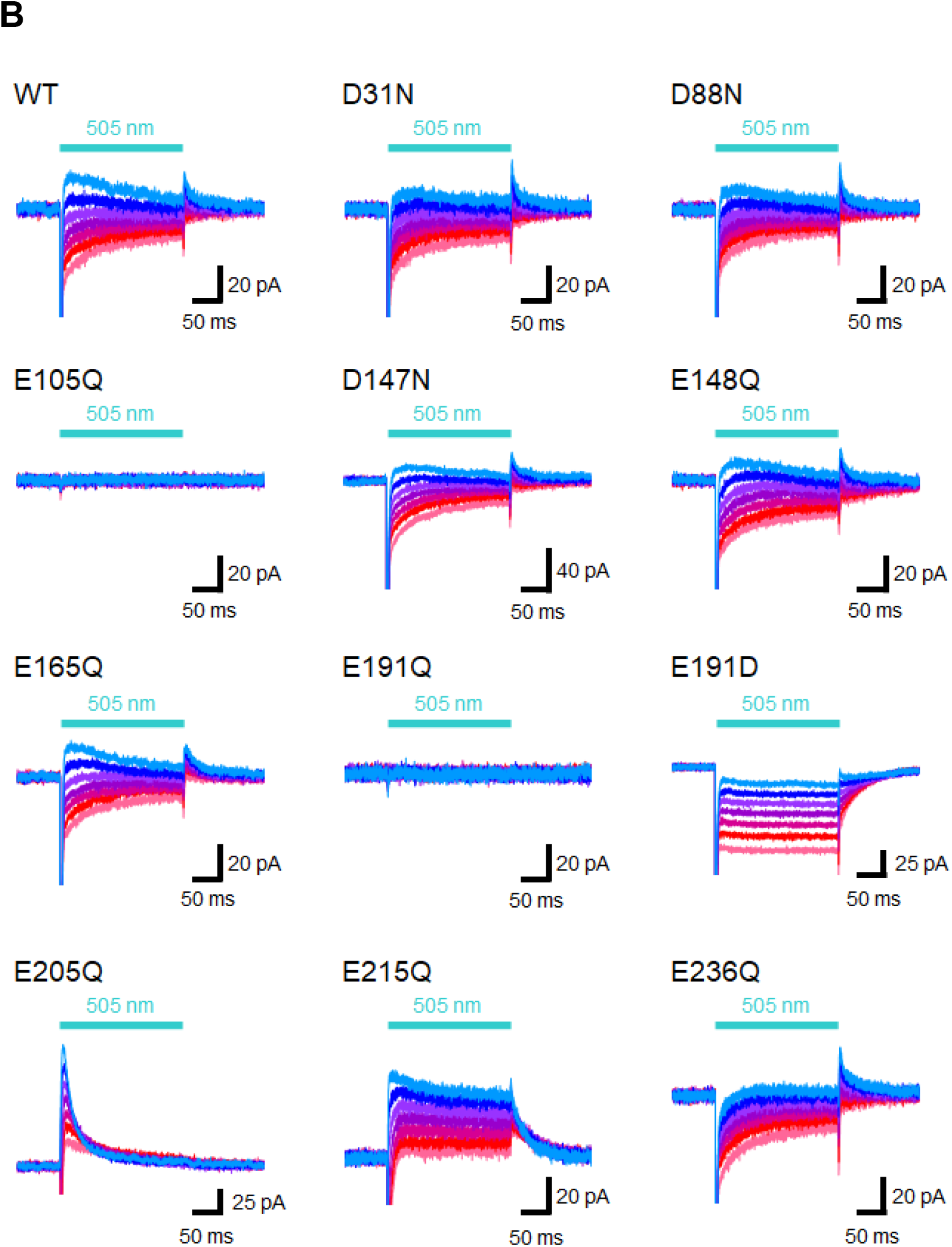
Representative photocurrent traces of the wild-type and mutantts of V2HeR3. (A) Electrophysiological measurements of V2HeR3 or variants-driven photocurrent in ND7/23 cells. The cells were illuminated with light (λ = 505 nm, 24.5 mW/mm^2^). The membrane voltage was clamped from −60 to +60 mV for every 20-mV step. Horizontal bar showed 50 ms. The pipette solution was 110 mM NaCl, pHi 7.4, the bath solution was 140 mM NaCl, pHe 7.4. (B) Focused on peak photocurrents and steady-state photocurrents region.

**Fig. Figure S12.**
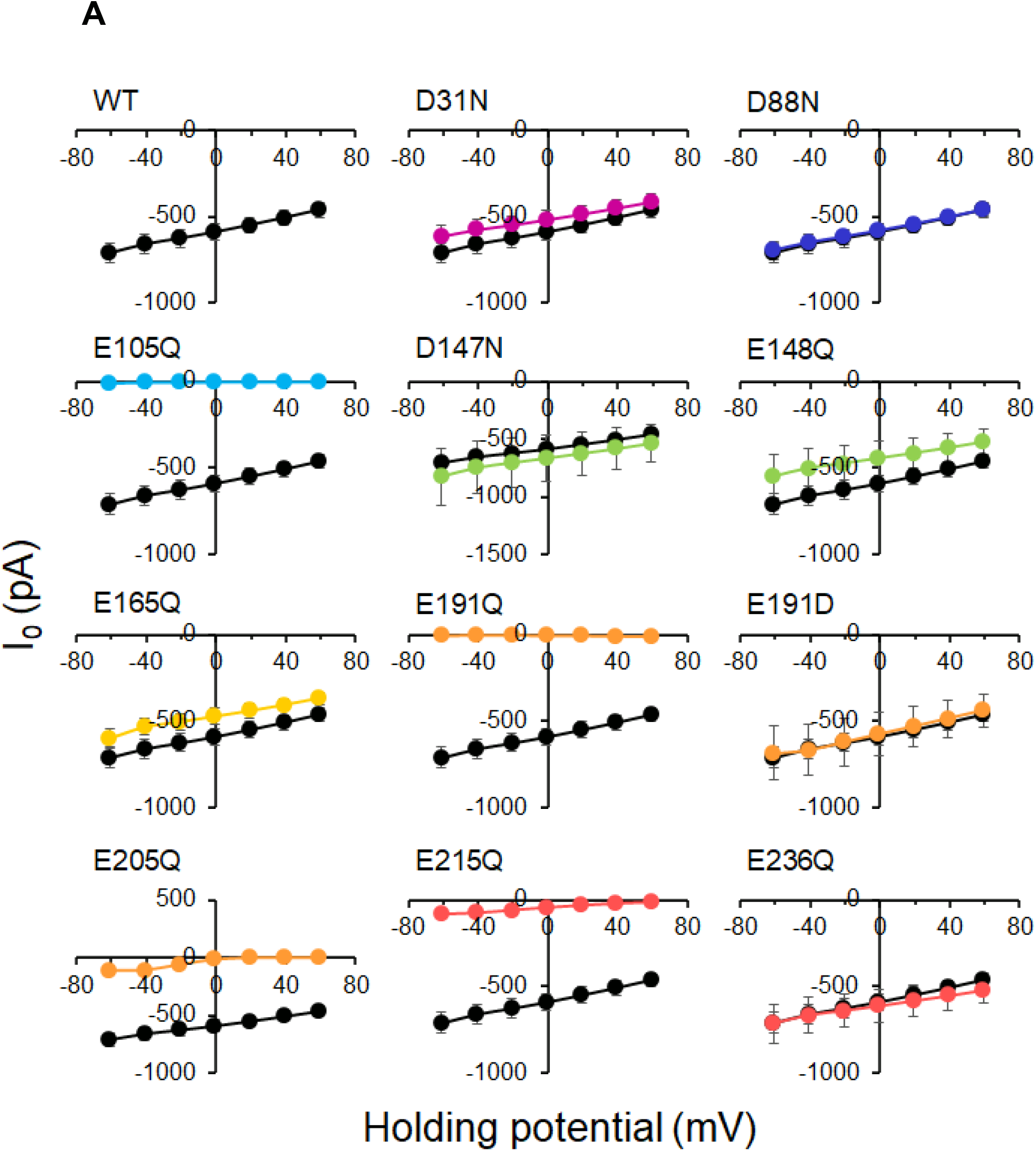
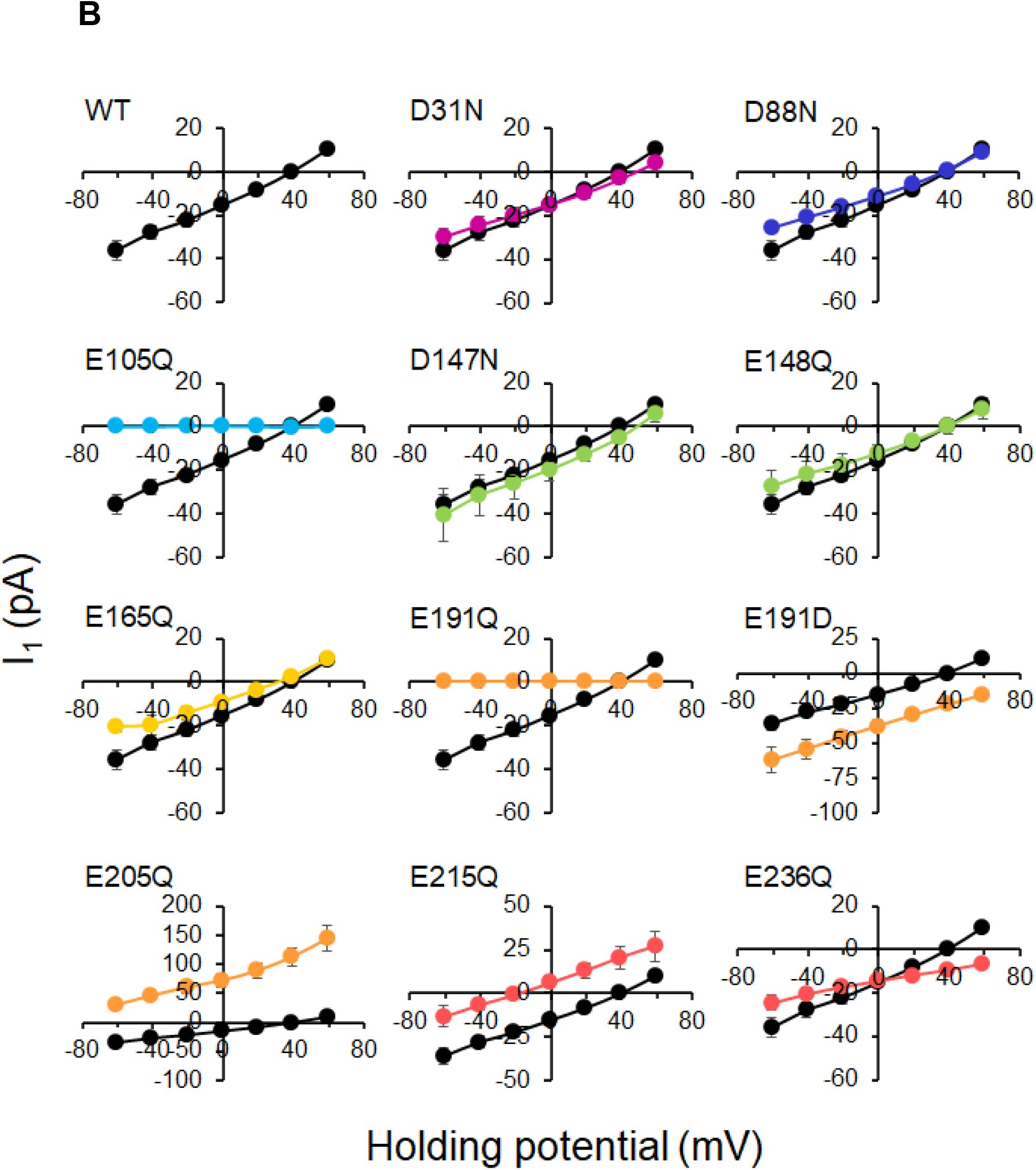
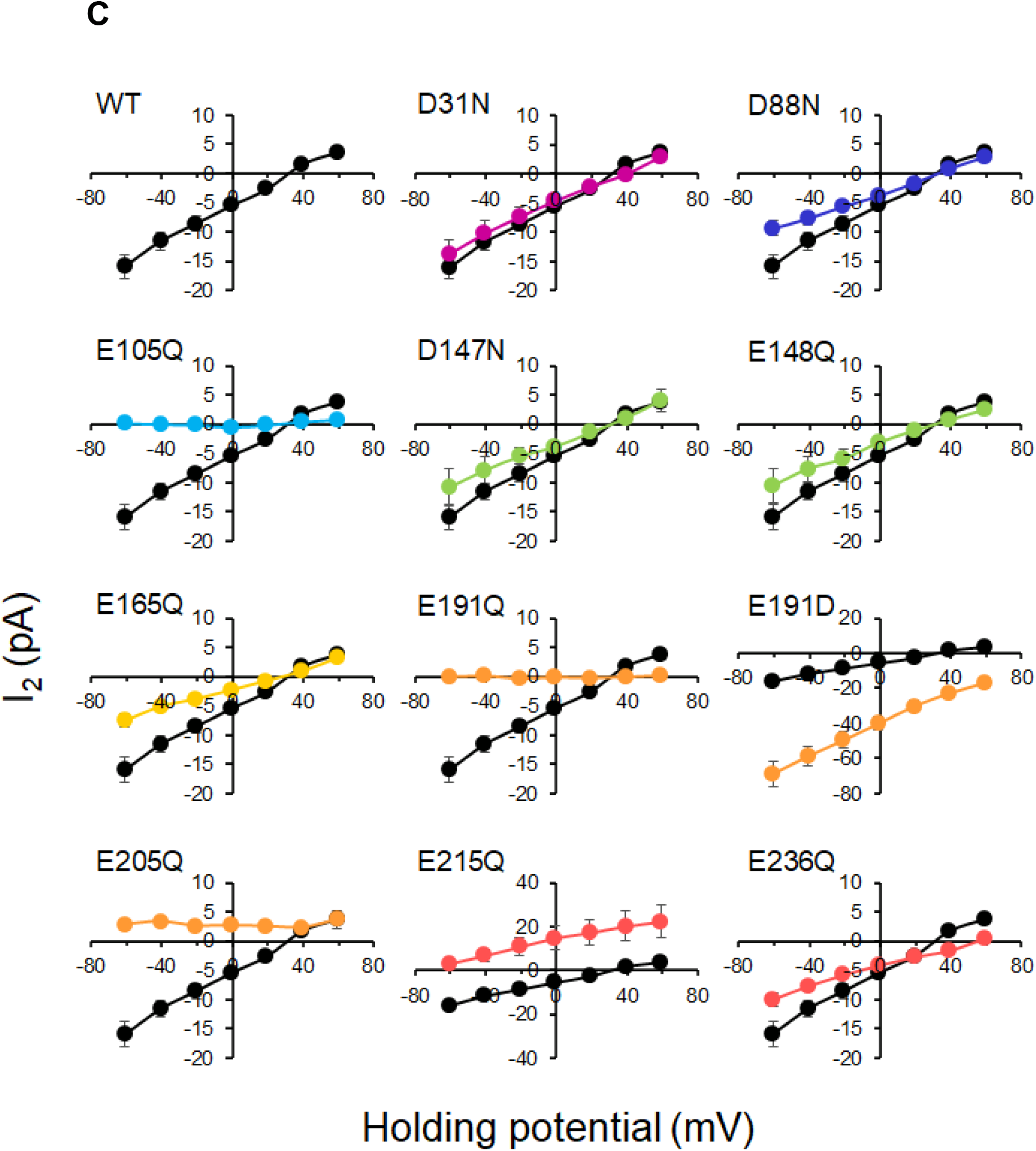
Photocurrent-Voltage plots of the wild-type and mutants of V2HeR3. Electrophysiological measurements of V2HeR3 or variants-driven photocurrent in ND7/23 cells. The cells were illuminated with light (λ = 505 nm, 24.5 mW/mm^2^). The membrane voltage was clamped from −60 to +60 mV for every 20-mV step. Black curves are the results of V2HeR3. The pipette solution was 110 mM NaCl, pHi 7.4, the bath solution was 140 mM NaCl, pHe 7.4. (A) Transient photocurrent-Voltage plots. (B) Peak photocurrent-Voltage plots. (C) Steady-state photocurrent-Voltage plots. n = 6 to 7 cells.

**Fig. Figure S13.**
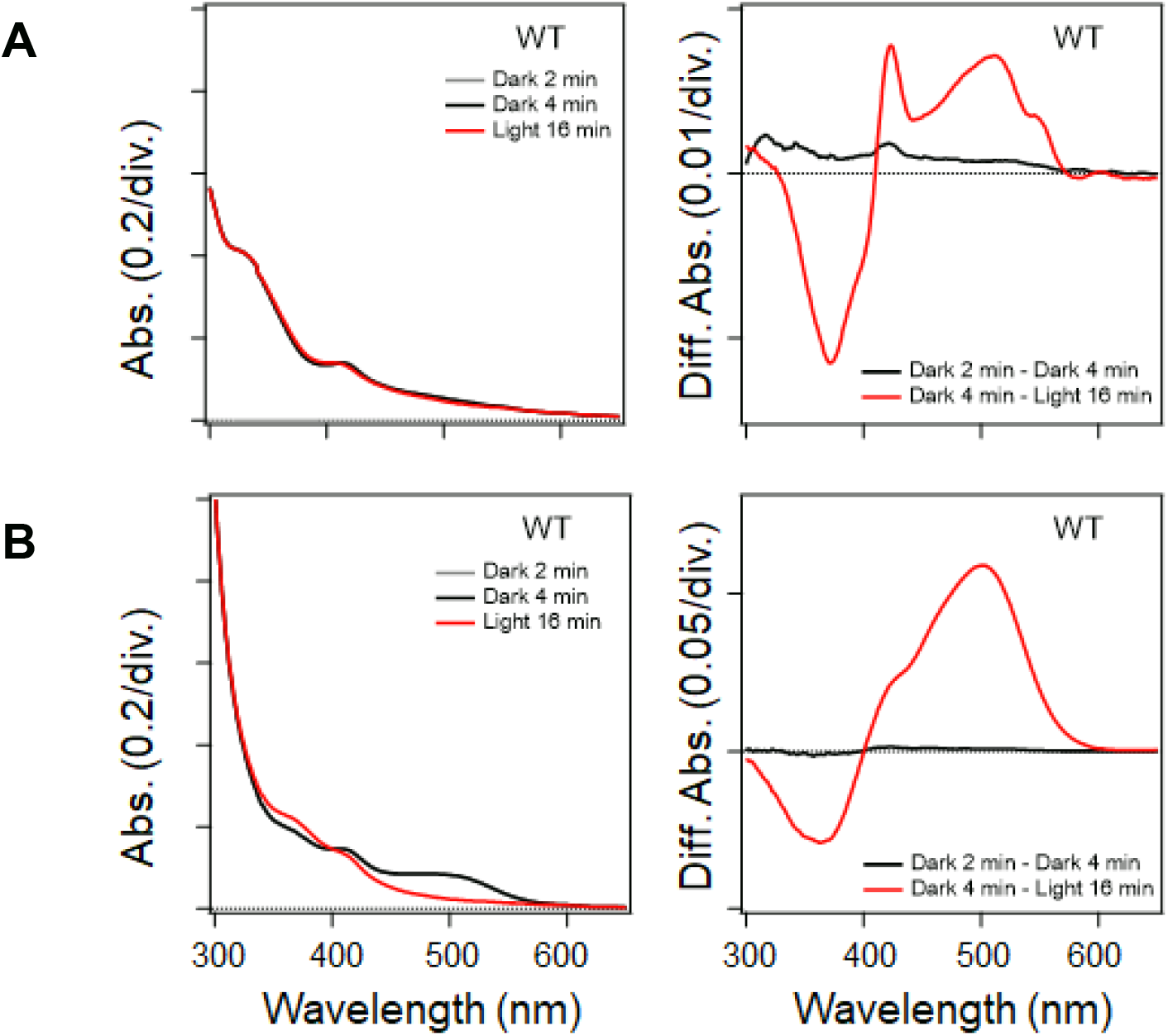
Absolute and difference absorption spectra of V2HeR3 in the experiments of hydroxylamine bleach. Light-induced difference absorption spectra in the presence of hydroxylamine. Left, Absolute absorption of DDM solubilized sample. Right, Difference absorption of before and after illumination (A) ND7/23 cells (B) *P. pastris* cells.

**Fig. Figure S14.**
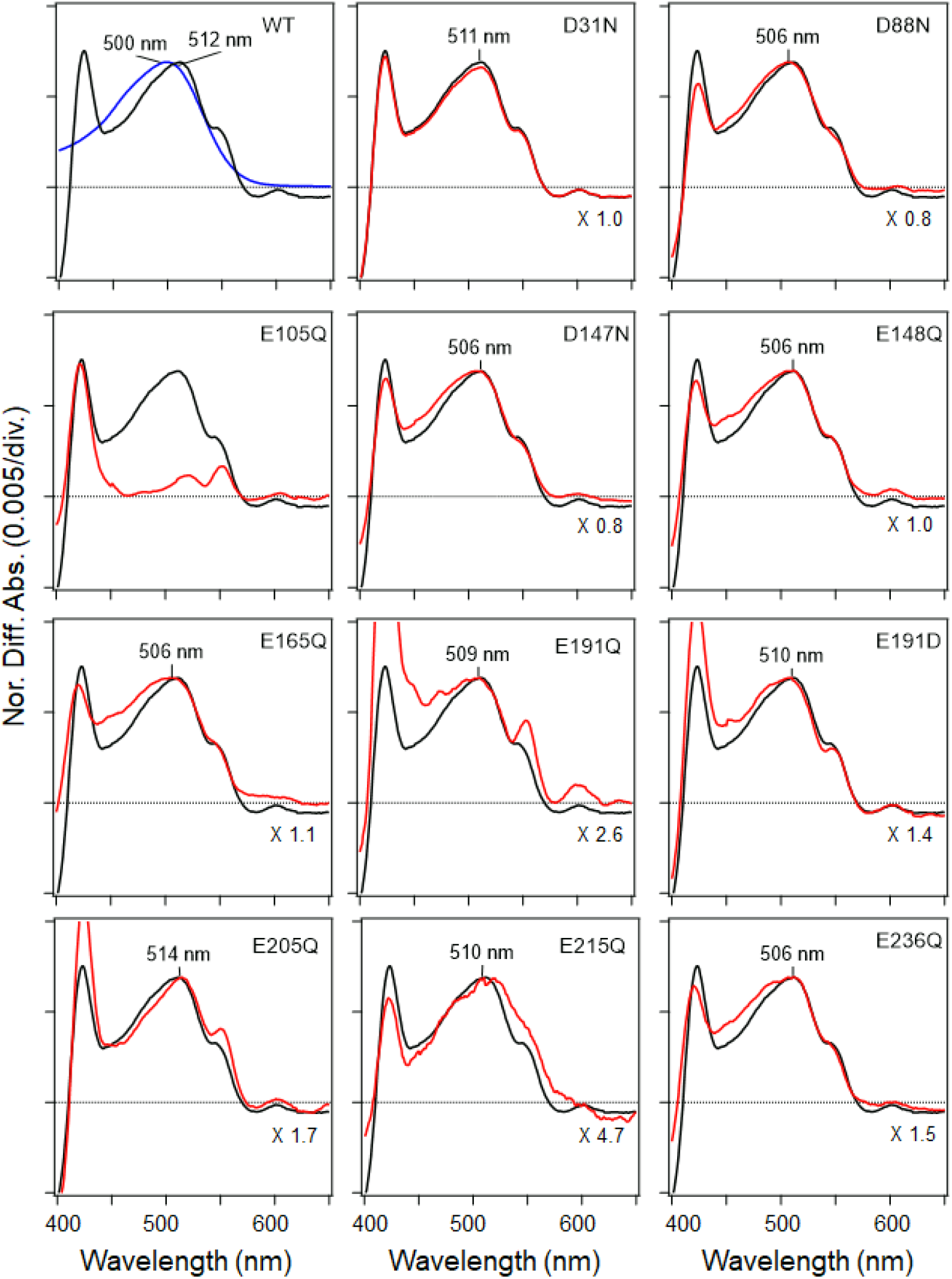
Absorption spectra of the wild-type and mutants of V2HeR3 obtained by the hydroxylxmine bleach of ND7/23 cells. Light-induced difference absorption spectra in the presence of 50 mM hydroxyl amine of the WT (black line) and the mutants (red line). Mutants and WT spectra were normalized by use of a positive peak at each λ_max_. Blue line in WT showed the absorption spectra of the purified V2HeR3.

**Fig. Figure S15.**
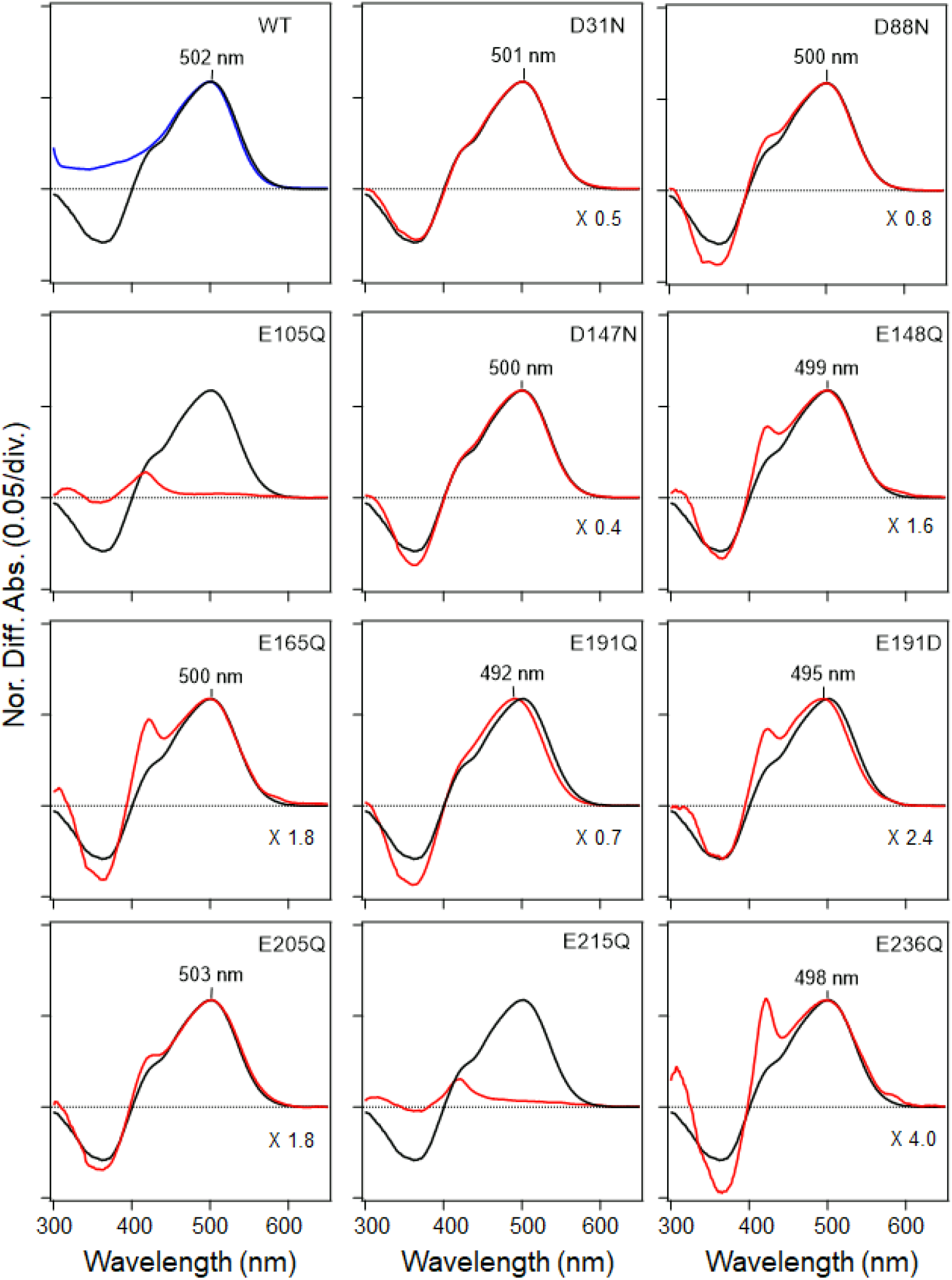
Absorption spectra of the wild-type and mutants of V2HeR3 obtained by the hydroxylxmine bleach of *Pichia pastris* cells. Light-induced difference absorption spectra in the presence of 0.5 M hydroxyl amine of the WT (black line) and the mutants (red line). Mutants and WT spectra were normalized by use of a positive peak at each λ_max_. Blue line in WT showed the absorption spectra of the purified V2HeR3.

**Fig. Figure S16.**
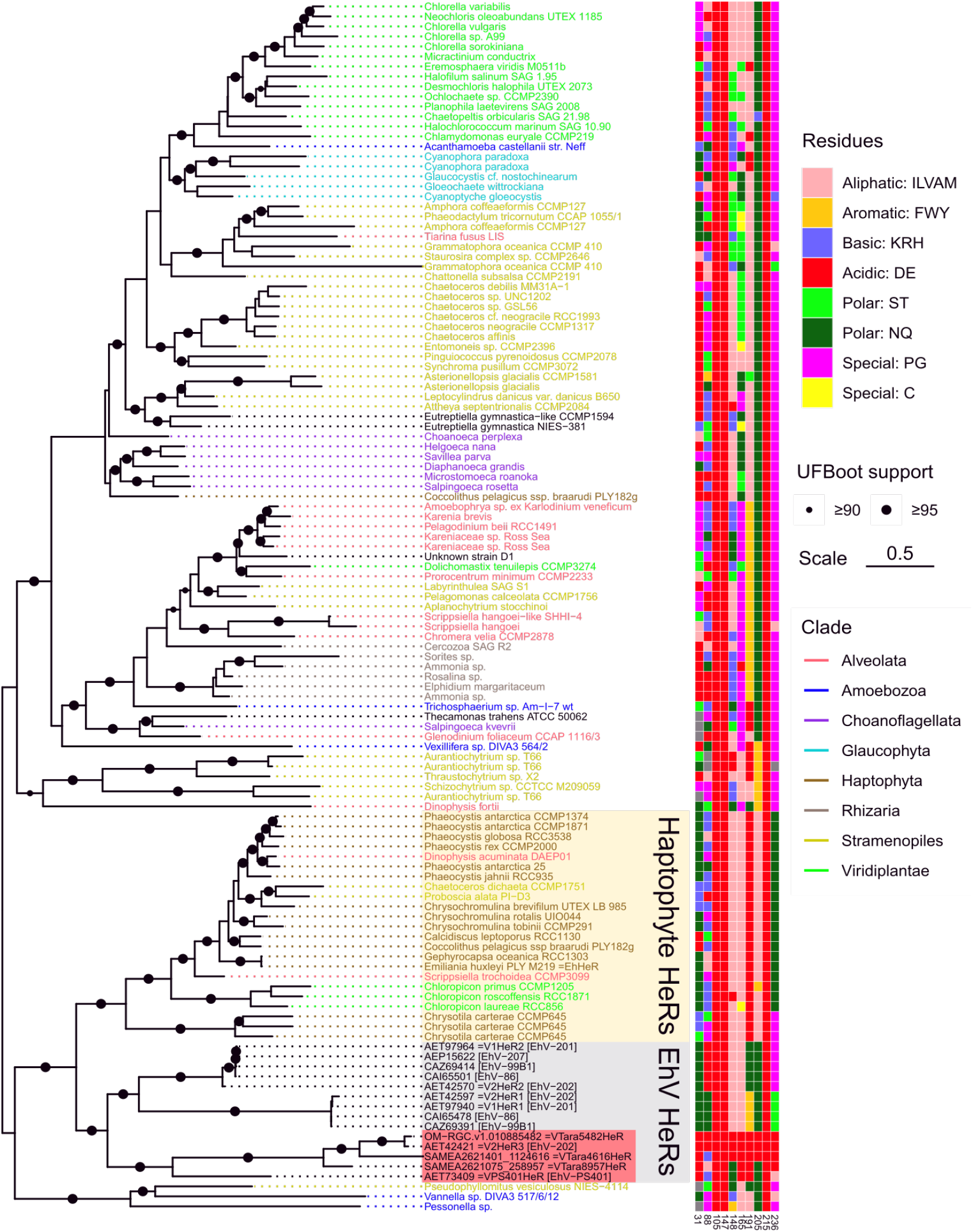
Phylogenetic relationships of the *Eh*V HeRs and their eukaryotic cognates. Eukaryotic HeRs most similar to HeRs from *Eh*V were collected from a collection of transcriptomes and genomes of algae and other unicellular eukaryotes. Eukaryotic HeRs are named after the corresponding species/strains and colored by their taxonomic origin. The tree is midpoint-rooted which places HeRs from *Eh*Vs as a sister clade to the predominantly haptophyte HeRs, a group that also includes *Eh*HeR from the host species, *E. huxleyi.* Distribution of the amino acid residues among a selected set of V2HeR3 positions is indicated to the right. Data and code used to generate the phylogeny are available from https://github.com/BejaLab/ViralHeRs.

**Fig. Figure S17.**
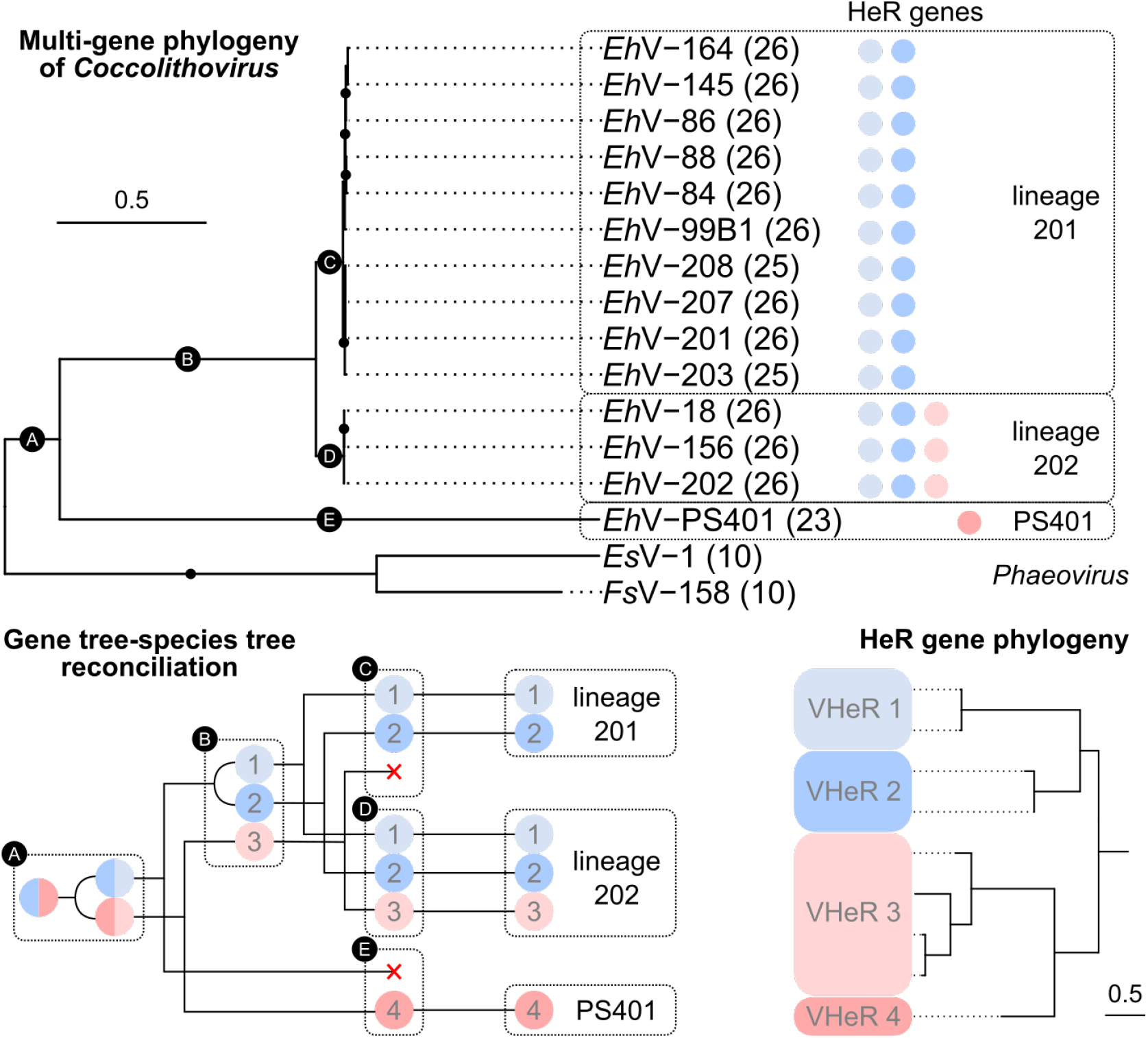
Evolution of coccolithoviruses and their HeR genes. Multigene phylogeny of *Eh*V isolates using 26 single-copy genes (number of orthogroups used per genome is indicated in parentheses) was rooted using two genomes from of phaeoviruses (top). Black dots represent highly supported branches (ultrafast bootstrap support ≥95). For each *Eh*V genome the presence of HeR genes from the four HeR clades is indicated with circles colored according to the HeR phylogeny (bottom right). The two clades showing proton transport are highlighted in red. Gene tree-species tree reconciliation providing a hypothesis about the evolution of the HeR genes among *Eh*Vs (bottom left) is based on the assumption of no horizontal transfer of HeR genes between viruses. ⊏ forks represent co-divergence, ⊂ forks represent gene duplication events, × represents gene losses. The numbers in the black circles correspond to branches on the species tree. List of phylogenetic markers, accession numbers of HeR genes and lineage assignments are available in Dataset S2.

**Fig. Figure S18.**
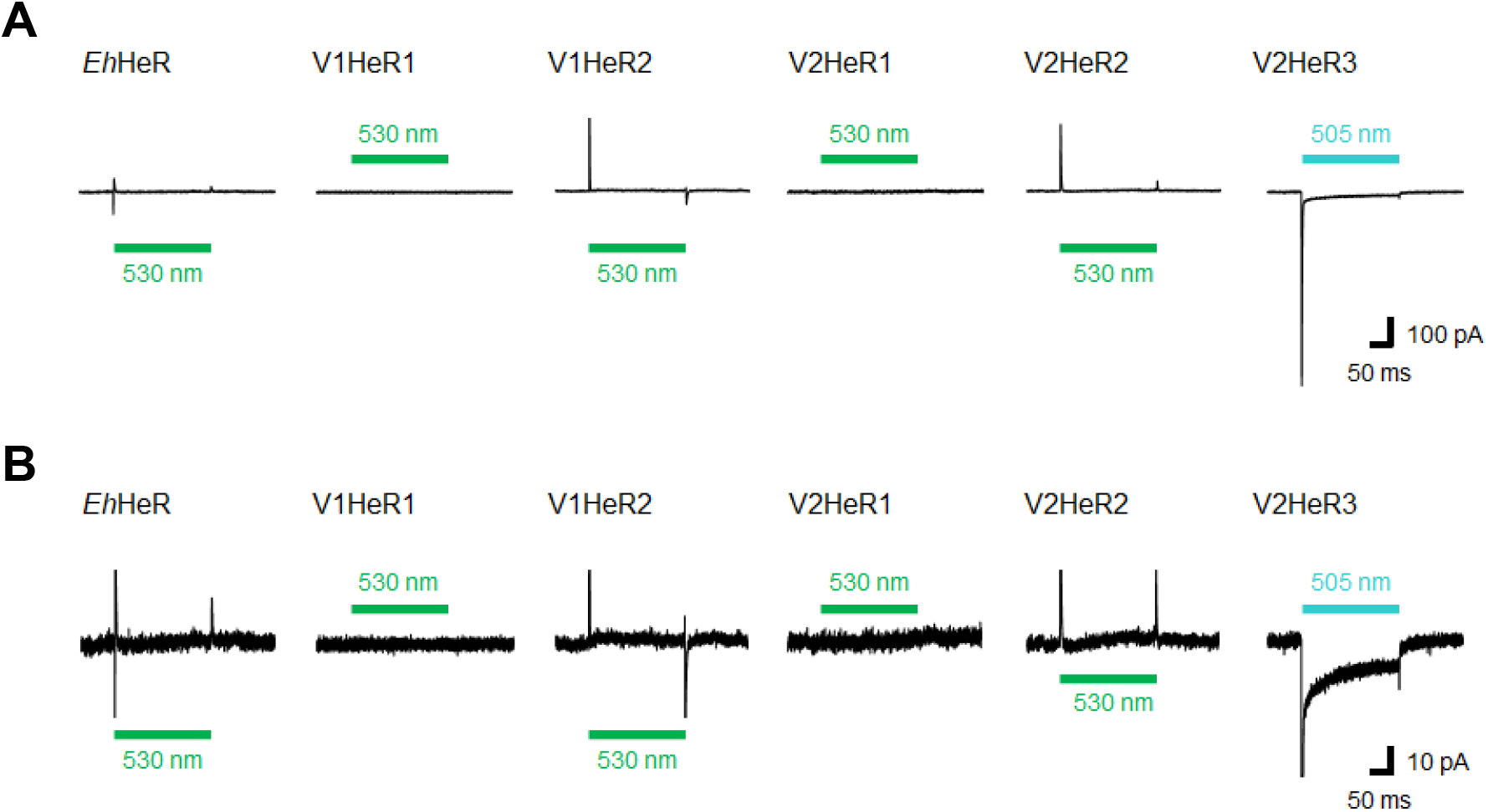
Representative photocurrent traces of HeRs from *E. huxleyi* and its viruses. (A) Electrophysiological measurements of virus HeRs driven photocurrent in ND7/23 cells. The cells were illuminated with light (λ = 505 nm, 24.5 mW/mm^2^, λ = 530 nm, 27.8 mW/mm^2^) during the time region shown by green bars. The membrane voltage was clamped −40 mV. The pipette solution was 110 mM NaCl, pHi 7.4, the bath solution was 140 mM NaCl, pHe 7.4. (B) Focused on peak photocurrents and steady-state photocurrents region.

**Fig. Figure S19.**
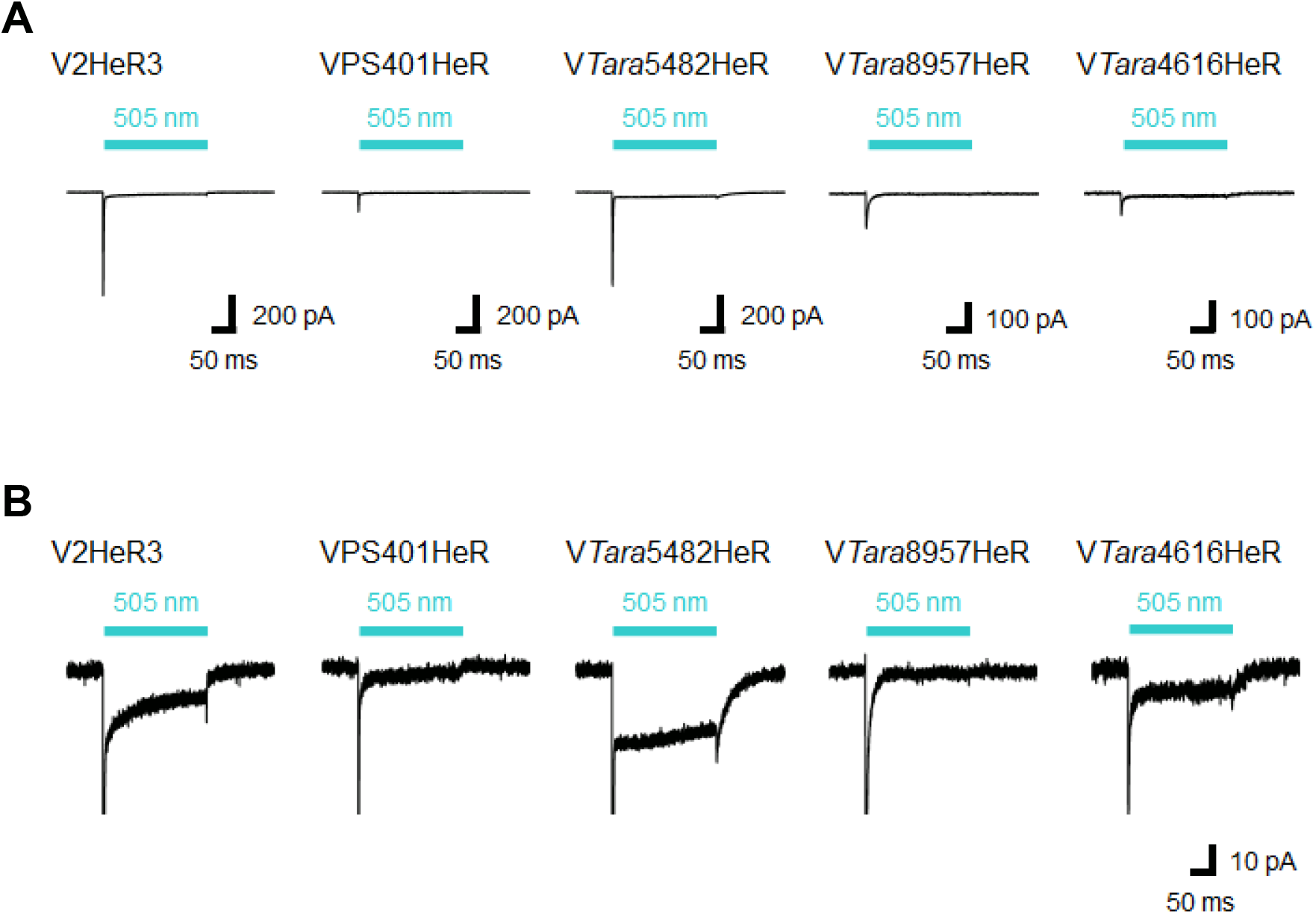
Representative photocurrent traces of V2HeR3-like HeRs. (A) Electrophysiological measurements of V2HeR3-like HeRs driven photocurrent in ND7/23 cells. The cells were illuminated with light (λ = 505 nm, 24.5 mW/mm^2^) during the time region shown by blue bars. The membrane voltage was clamped −40 mV. The pipette solution was 110 mM NaCl, pHi 7.4, the bath solution was 140 mM NaCl, pHe 7.4. (B) Focused on peak photocurrents and steady-state photocurrents region.

**Fig. Figure S20.**
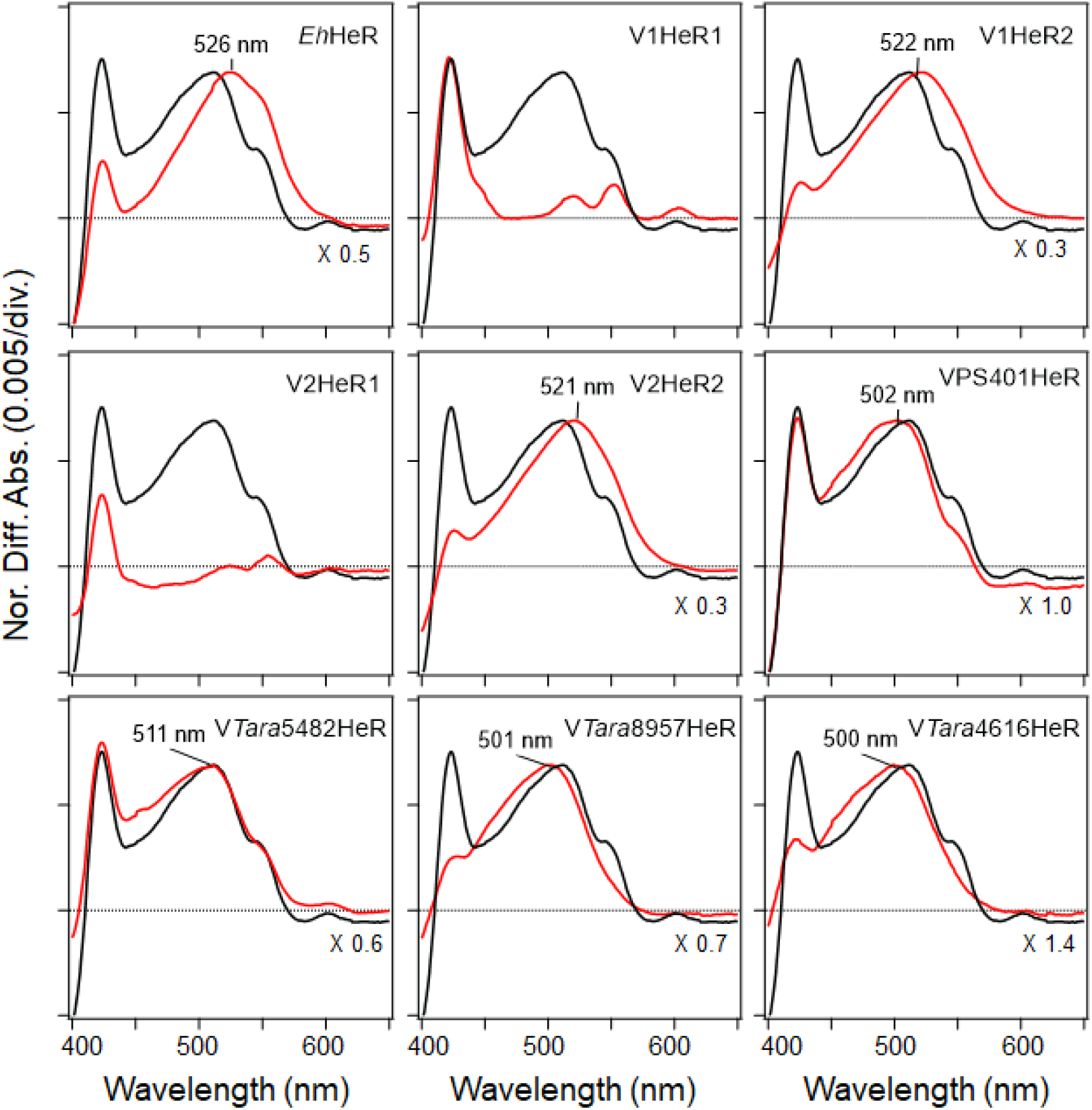
Absorption spectra of HeRs from *E. huxleyi* and its viruses, and V2HeR3-like HeRs obtained by the hydroxylxmine bleach of ND7/23 cells. Light-induced difference absorption spectra in the presence of 50 mM hydroxyl amine of the V2HeR3 (black line) and the *E. huxleyi* and its viruses, and V2HeR3-like HeRs (red line). Absorption spectra were normalized by use of a positive peak at each λ_max_.

**Fig. Figure S21.**
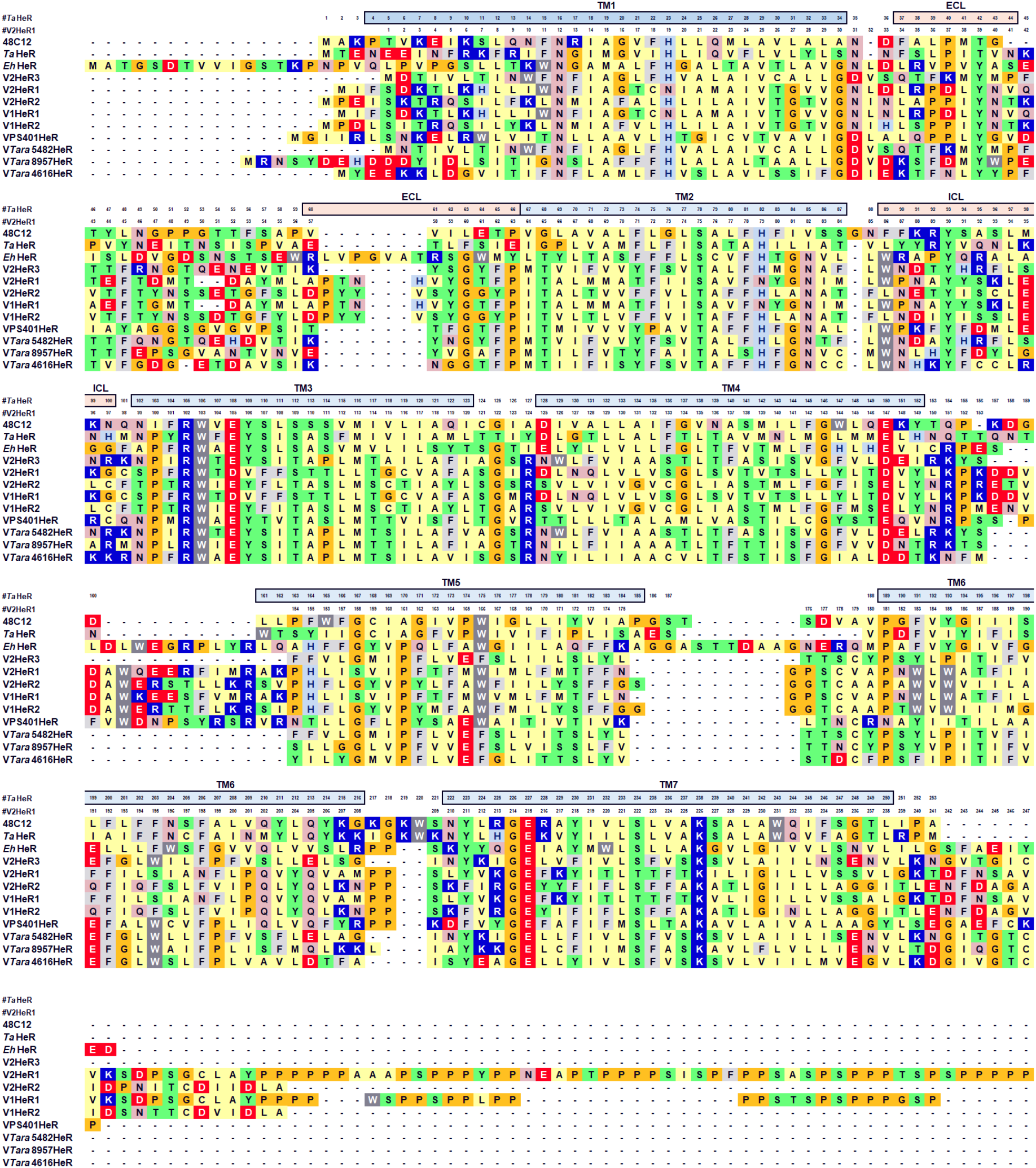
Alignment of HeRs. Multiple amino acid alignment of V2HeR3 with 48C12, *Ta*HeR, *Eh*HeR, V1HeR1, V1HeR2, V2HeR1. V2HeR2, VPS401HeR, V*Tara*5482HeR, V*Tara*4616HeR and V*Tara*8957HeR.

**Table S1.** Pairwise protein sequence identities (%) between the tested HeRs in the transmembrane regions (total alignment length of 150 residues).

|  | VPS401<br>HeR | V2HeR3 | VTara5482<br>HeR | VTara4616<br>HeR | VTara8957<br>HeR | V2HeR2 | V1HeR2 | V2HeR1 | V1HeR1 | EhHeR |
| --- | --- | --- | --- | --- | --- | --- | --- | --- | --- | --- |
| VPS401HeR |  | 36.7 | 35.3 | 37.3 | 40.7 | 38.7 | 38 | 22 | 22 | 31.3 |
| V2HeR3 | 36.7 |  | 92.7 | 64.7 | 62.7 | 35.3 | 35.3 | 27.3 | 27.3 | 28.7 |
| VTara5482HeR | 35.3 | 92.7 |  | 64.7 | 61.3 | 34 | 34 | 27.3 | 26.7 | 28.7 |
| VTara4616HeR | 37.3 | 64.7 | 64.7 |  | 58.7 | 34.7 | 34 | 23.3 | 23.3 | 33.3 |
| VTara8957HeR | 40.7 | 62.7 | 61.3 | 58.7 |  | 34 | 35.3 | 22.7 | 22.7 | 28.7 |
| V2HeR2 | 38.7 | 35.3 | 34 | 34.7 | 34 |  | 90 | 32.7 | 32.7 | 42 |
| V1HeR2 | 38 | 35.3 | 34 | 34 | 35.3 | 90 |  | 32.7 | 32.7 | 39.3 |
| V2HeR1 | 22 | 27.3 | 27.3 | 23.3 | 22.7 | 32.7 | 32.7 |  | 96 | 26 |
| V1HeR1 | 22 | 27.3 | 26.7 | 23.3 | 22.7 | 32.7 | 32.7 | 96 |  | 24.7 |
| EhHeR | 31.3 | 28.7 | 28.7 | 33.3 | 28.7 | 42 | 39.3 | 26 | 24.7 |  |

**Table S2.** Genes encoded on the metagenomic sequences encoding HeR genes related to V2HeR3 (AET42421.1). Best-matching blastp hits against the nr database are provided.

| CDS | Function | Accession | Locus tag | Organism | Score | Query coverage | E-value | Identity |
| --- | --- | --- | --- | --- | --- | --- | --- | --- |
| Contig SAMEA2621075_258957 |  |  |  |  |  |  |  |  |
| tran_2 | Heliorhodopsin | AET42421.1 | EXVG_00394 | EhV-202 | 284 | 96% | 4.00E-93 | 57.90% |
| Contig SAMEA2621401_1124616 |  |  |  |  |  |  |  |  |
| tran_2 | Heliorhodopsin | AET42421.1 | EXVG_00394 | EhV-202 | 287 | 95% | 3.00E-94 | 57.50% |
| tran_3 | Thioredoxin | AEO98306.1 | ELVG_00005 | EhV-203 | 151 | 76% | 4.00E-42 | 43.70% |
| Contig TARA_B100000767_G_C19749265_1 |  |  |  |  |  |  |  |  |
| tran_3 | Heliorhodopsin | AET42421.1 | EXVG_00394 | EhV-202 | 464 | 100% | 3.00E-164 | 91.50% |
| tran_4 | Unknown | AET42406.1 | EXVG_00446 | EhV-202 | 219 | 100% | 1.00E-71 | 89.60% |
| tran_5 | Unknown | AET42405.1 | EXVG_00445 | EhV-202 | 108 | 81% | 1.00E-28 | 75.00% |

**Table S3.** Incidence of the *Eh*HeR gene among *Emiliania huxleyi* transcriptomes. Matches were obtained by using blastp against the NCBI TSA database. Note that the gene is absent from the genome assembly of *Emiliania huxleyi* CCMP1516 (NCBI Assembly GCA_000372725.1).

| Strain | NCBI TSA contigs |
| --- | --- |
| CCMP3266 | GIZZ01039194.1, GIZZ01031796.1, GIZZ01022151.1, GIZZ01020795.1 |
| CCMP370 | HBOC01014818.1, HBOC01014817.1, HBOC01014816.1, HBOC01014815.1, HBOC01014814.1, HBOC01014813.1 |
| CCMP371 | GHJP01012634.1 |
| PLY M219 | HBOB01045754.1, HBOB01045753.1, HBOB01045752.1, HBOB01045751.1, HBOB01045750.1, HBOB01045749.1, HBOB01045748.1, HBOB01045747.1, HBOB01045746.1, HBOB01045745.1, HBOB01045744.1, HBOB01045743.1 |
| RCC1229 | HBTT01000153.1, HBTT01000152.1 |
| RCC1824 | HBTF01056801.1, HBTF01054040.1, HBTF01054039.1 |
| RCC912 | HBTW01055061.1, HBTW01038897.1, HBTW01038896.1 |

**Table S4.**
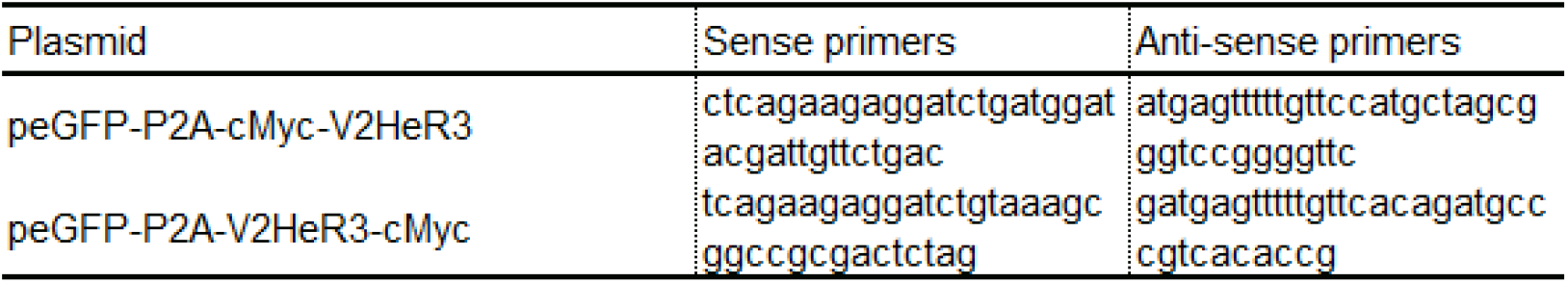
List of primers used for insertions.

**Table S5.**
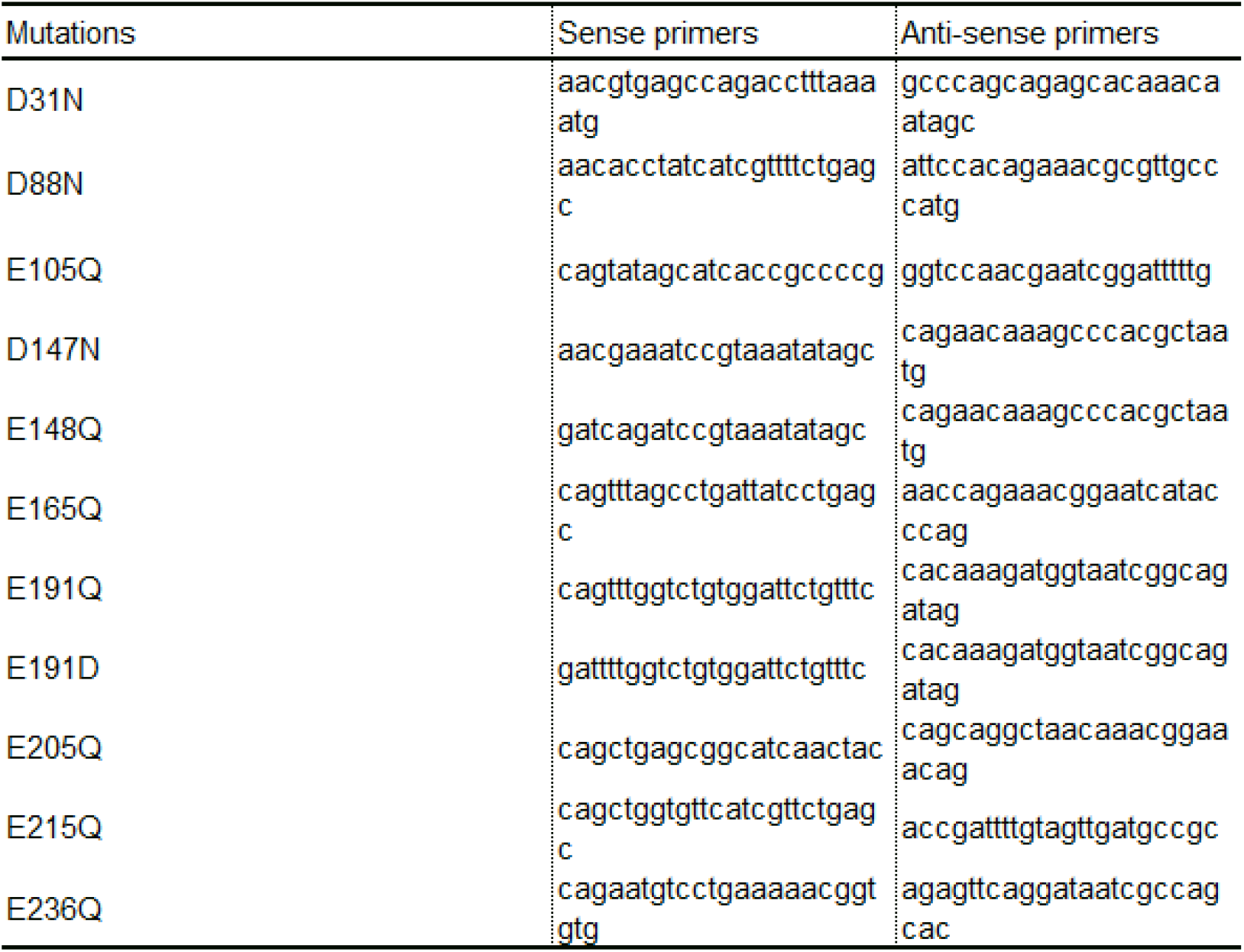
List of primers used for site-directed mutagenesis.

**Table S6.**
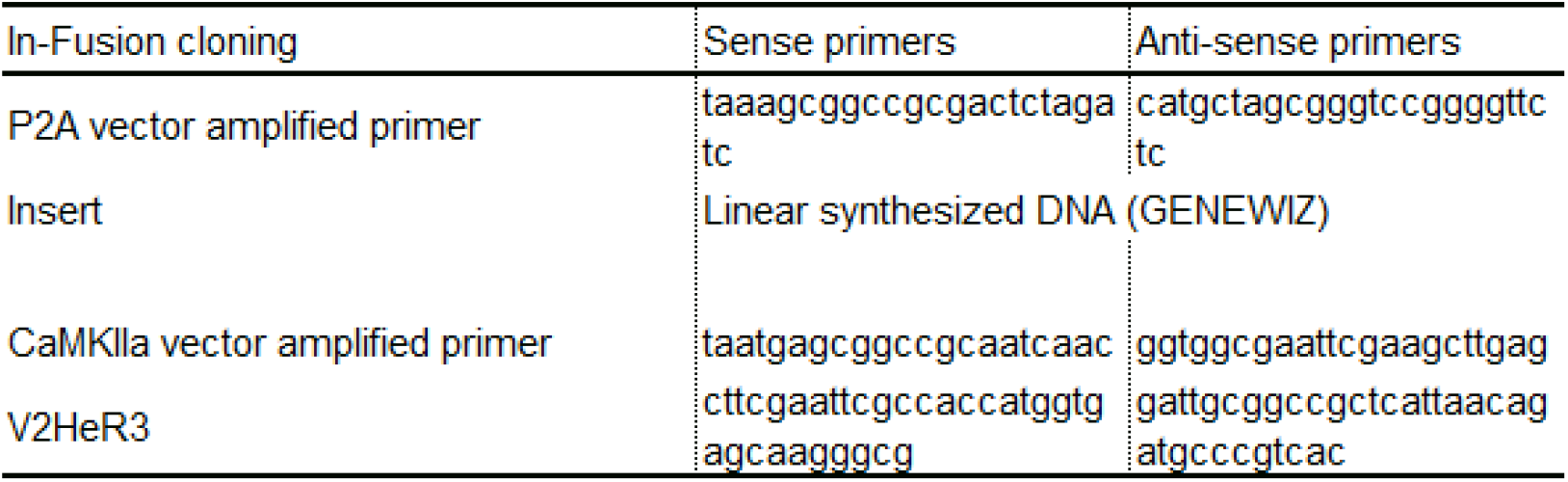
List of primers used for subcloning into an eGFP-P2A vector.

**Table S7.**
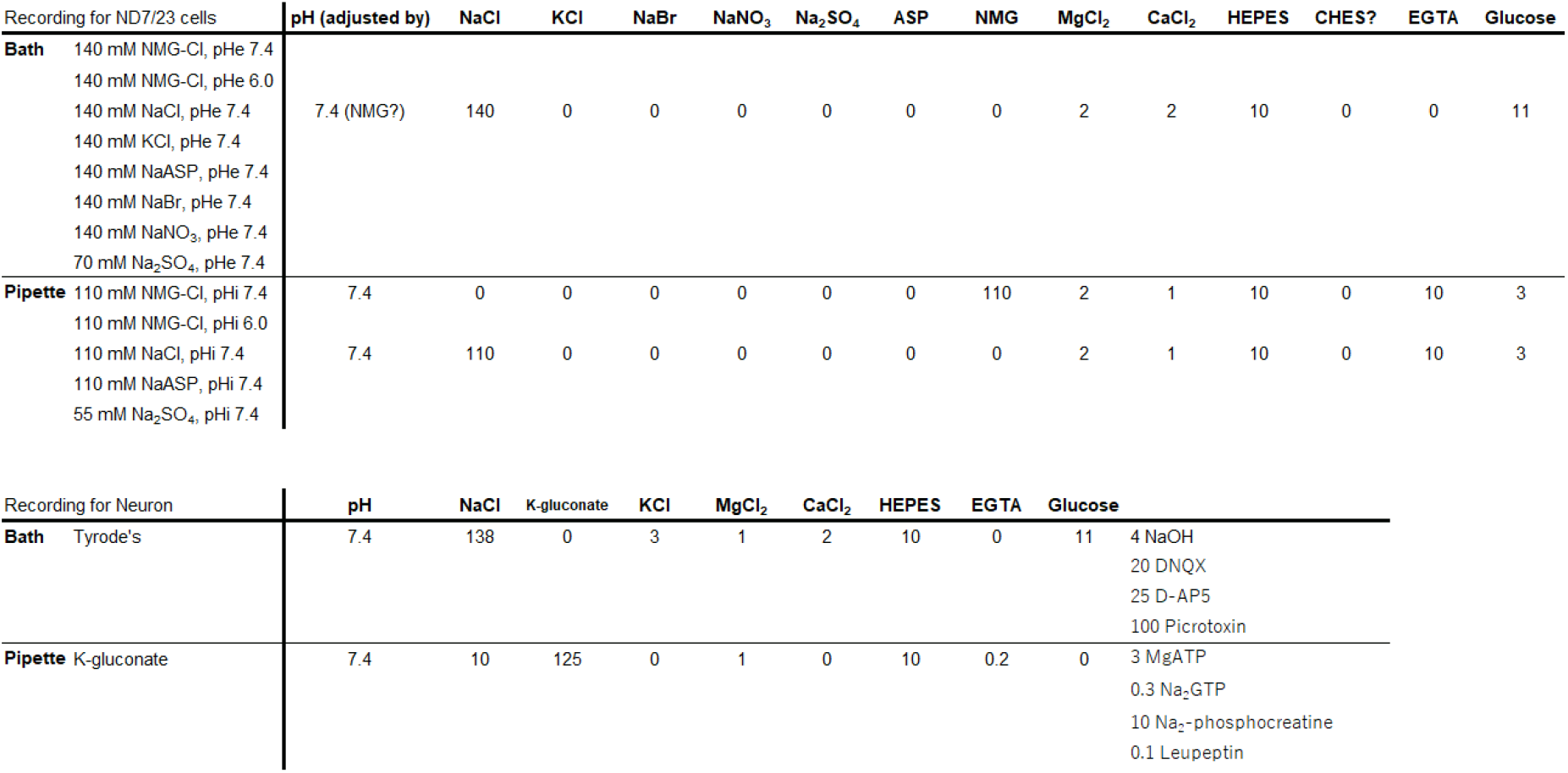
Composition of pipette and bath solutions in whole-cell voltage or current clamp. Abbreviations: Asp, aspartate; NMG, N-Methyl-D-glucamine; HEPES, 4-(2-hydroxyethyl)-1-piperazineethanesulfonic acid; EGTA, ethylene glycol tetraacetic acid; DNQX, 6,7-Dinitroquinoxaline-2,3-dione; D-AP5, D-(−)-2-amino-5-phosphonopentanoic acid. All concentrations are in mM.

**Dataset S1 (Genbank flat file).** Annotated sequences of the metagenomic contigs containing V*Tara*5482HeR, V*Tara*8957HeR and V*Tara*4616HeR.

**Dataset S2 (Excel spreadsheet).** List of viral genomes and phylogenetic markers used in this study. For each virus, accessions of HeRs genes and accessions for the indicated phylogenetic markers are provided.

